# Multi trait assessment of wheat variety mixtures performance and stability: mixtures for the win!

**DOI:** 10.1101/2024.07.22.604587

**Authors:** Laura Stefan, Silvan Strebel, Karl-Heinz Camp, Sarah Christinat, Dario Fossati, Christian Städeli, Lilia Levy Häner

## Abstract

In the current quest for a more sustainable, environment-friendly agriculture, variety mixtures are often suggested as a practical option to increase the stability of food production systems. Their effects on yield have been extensively researched, yet clear conclusions remain elusive, notably in terms of mechanistic processes and optimal variety combinations. Furthermore, in the case of wheat, yield is not the only component in the equation: grain quality is crucial for the bread value chain, yet the effects of variety mixtures on wheat quality and its stability have rarely been investigated. To that end, we conducted a multi-year, multi-site wheat variety mixture experiment investigating the role of variety mixtures on the performance and stability of five traits linked to grain yield and quality, and the mechanisms underlying these effects. Eight varieties were grown in pure stands and 2-variety mixtures, following a full diallel design. We considered the responses of grain yield, protein content, thousand kernel weight, hectoliter weight, and Zeleny sedimentation value. Results showed that mixtures generally outperformed pure stands in terms of global performance and stability for the 5 parameters. We particularly noticed an increase in quality stability and in Zeleny sedimentation value in mixtures, showing the potential of mixtures to improve crop quality. Moreover, we highlighted the important role of light interception for increased mixtures benefits. A more detailed investigation into individual mixture performances led us to some practical rules for optimal variety combinations: we advise combining varieties with similar heights and phenologies but different tillering abilities and yield potential. This study thus shows that variety mixtures represent a promising solution to sustainably increase the stability of wheat yield and quality. With practical recommendations, our results could benefit farmers but also processors and bakers, and promote the adoption of wheat variety mixtures.

## Introduction

Food production systems will face increasingly challenging conditions in the near future (Tilman et al., 2011). Growing human population will require increasing, or at least maintaining crop yields (van Dijk et al., 2021). At the same time, climate change is expected to increase the frequency and magnitude of extreme weather events, such as drought, heat, or heavy rainfall, which can greatly impact agricultural systems (IPCC, 2021; Jägermeyr et al., 2021). On top of this, there is an urgent need to reduce the detrimental environmental impacts of crop production systems, through a reduction in synthetic inputs, such as mineral fertilizers, pesticides, and herbicides (Baweja et al., 2020; Thomas et al., 2020). It is thus crucial to design agricultural production systems that are at the same time equally or more performant, more stable, and less reliant on external inputs (Bommarco et al., 2013). Increasing diversity in agricultural systems appears as a potential solution (Gurr et al., 2016; Reckling et al., 2022; Tamburini et al., 2020), but increasing diversity at the species level comes with technical and processing challenges that may hinder its adoption (Brooker et al., 2015; Hong et al., 2020; Lithourgidis et al., 2011). Variety mixtures combine the best of the two worlds, since it allows to increase diversity at the genotype level while remaining similar as the standard culture in terms of machinery, technical culture, and equipment (Wuest et al., 2021).

It has been shown in the literature that variety mixtures often lead to benefits in yield production (Fletcher et al., 2019; Tooker & Frank, 2012). While present, these benefits remain small, with an average of 3-4% yield increase in mixtures compared to pure varieties according to large meta-analyses (Borg et al., 2017; Huang et al., 2024). Increasing productivity is not the only service delivered by variety mixtures: other benefits include decrease in pest and disease pressures (Kristoffersen et al., 2020; Vidal et al., 2020), decrease in weed pressure, potential increase in grain quality (Lazzaro et al., 2018), and potential increase in the stability of grain yield and quality (Döring et al., 2015; L. Li et al., 2023; Stefan et al., 2024). Despite decades of investigation, the mechanisms – and their importance – responsible for these mixture benefits remain unclear, which makes it difficult to choose which varieties to assemble (L. Li et al., 2023). For instance, some studies highlight the importance of varying susceptibilities to pest and diseases (Finckh et al., 2000; Kristoffersen et al., 2021), or the predominant role of asynchrony for stability (Stefan et al., 2024; Wuest et al., 2021), while some effects are thought to arise from complementarity between the varieties, leading to a reduction in competition for resources (Barot et al., 2017; Revilla-Molina et al., 2009). Notably, a few recent studies have emphasized the role of competition for light for grain yield overyielding: Tschurr et al. (2023) demonstrated that grain overyielding in oat variety mixtures was well explained by differences in canopy covers between mixtures and pure stands, suggesting that mixtures show higher potential for efficient light interception. In a study on wheat variety mixtures, Gawinowski et al. (2024) showed the importance of tillering plasticity due to shade avoidance – i.e. the ability of a variety to make more tillers in response to increased shade in mixtures – to explain overyielding. This provides first steps into mechanistic understanding of mixture yield benefits.

In addition to the uncertainties linked to the mechanisms, questions remain open regarding the context-dependence of mixture benefits. Ecological theory and experiments have indeed shown that plant-plant interactions strongly depend on the environmental context and harshness (Bertness & Callaway, 1994; Lortie & Callaway, 2006; Maestre et al., 2009), but the direction and magnitude of these shifts is less clear (Brooker et al., 2021; Maestre et al., 2006). It is often thought that species or variety mixtures are more beneficial in stressful conditions, as we then expect more facilitation and more compensation between the species/varieties (Darch et al., 2018; Steudel et al., 2012). This was notably demonstrated by Su et al. (2023) in the case of maize variety mixtures: in their study, variety mixtures stabilized aboveground biomass production only under stressful conditions (i.e. without irrigation, and with lower soil fertility). They also showed that complementarity effects were positive and facilitation more important at the stressful site, suggesting that increasing variety diversity is better suited under stressful and more fluctuating environmental conditions (Su et al., 2023). Other findings are in contradiction with this pattern (Alsabbagh et al., 2022): in some cases, the highest mixture benefits (such as overyielding) were found in highly productive environments (Chen et al., 2021; C. Li et al., 2020; Stefan, Engbersen, et al., 2021). Higher productivity can indeed lead to more competition between plants, and, therefore, higher benefits of species/variety complementarity – that is, reduced competition – in mixtures (Goldberg & Novoplansky, 1997; Hooper et al., 2005; Stefan et al., 2022). Shedding light on these environmental uncertainties is essential to provide accurate and adapted advice to farmers for efficient diversification of their crop production.

The bulk of research on variety mixtures has been focusing on the effects of variety diversity on grain crop yield, as it remains the most important parameter for farmers’ income (e.g. see (Borg et al., 2017; Döring et al., 2015; Kiær et al., 2009). However, grain yield alone does not give a complete image of the agronomic performance of a crop. In addition to yield, grain quality needs to be evaluated in order to fully assess crop performance (Levy Häner et al., 2015; Matzen et al., 2019; Reddy et al., 2003). Among quality traits, protein content and composition are essential, as proteins are crucial food components, vital for human and animal health (Fox et al., 1992; Wu, 2016). In the case of cereals, such as wheat, proteins include different types of glutens, which directly impact the quality and taste of the final product (e.g. bread) (Schopf et al., 2021; Wieser et al., 2023). Besides protein content, it is common in the wheat processing industry to measure Zeleny sedimentation value, which gives insights into gluten content and quality (Hrušková & Faměra, 2003; Levy Häner et al., 2015). Other crop quality traits include kernel weight, often measured as Thousand Kernel Weight (TKW), and specific weight, often measured as Hectoliter Weight (HLW). These parameters are important indicators of the physical quality of grains and are commonly recognized as indicators of potential flour yield by the industry (Marconi et al., 1999; Sirat, 2023). Additionally, the test weight has (together with protein content) a direct influence on the price the farmer receives for his wheat grains (swissgranum, 2023a). To the best of our knowledge, very few studies have investigated the effects of variety mixtures on the performance of both crop yield and crop quality parameters (Alsabbagh et al., 2022; Hoang et al., 2021; Lazzaro et al., 2018; Mille et al., 2006), and even fewer have additionally looked at the stability of several quality parameters (Tremmel-Bede et al., 2016).

To fill this knowledge gap, we investigated the performance and stability of wheat variety mixtures in a comprehensive way, by assessing not only crop grain yield, but also several key parameters linked to crop quality. Wheat was chosen in this study as it represents the third major grain crop worldwide in terms of production (FAO, 2022) and accounts for 29.8% of the arable land in Switzerland, which furthermore possesses a strong wheat breeding program (Federal Statistical Office, 2022). From 2020 to 2023, we set up a large variety mixture field experiment involving 8 Swiss varieties candidates, that were sown either in pure stands, in every possible 2-variety mixture (full diallel design), and in 8-varieties mixtures. The experiment was repeated in 3 sites across Switzerland for 3 growing seasons to evaluate stability of the mixtures. We measured crop grain yield to assess productivity, and grain protein content, TKW, HLW, and Zeleny sedimentation value to assess crop agronomic quality. Our research questions are the following: 1) are wheat variety mixtures better than pure stands in terms of agronomic performance and stability? 2) are performance and stability of mixtures context-dependent? 3) can we find general rules to optimise variety choices for increased performance and stability of mixtures? We expected variety mixtures to outperform pure stands, notably in terms of stability. We further expected that the benefits of mixtures for performance and stability would be higher in more stressful environments.

## Materials & Methods

### Experimental sites

The study took place over the course of three growing seasons – 2020/2021, 2021/2022 and 2022/2023 – in three sites across the Swiss Central Plateau. In the following, we will designate each growing season by its harvesting year (i.e. 2020/2021 will be referred to as 2021, 2021/2022 as 2022, and 2022/2023 as 2023). The experimental sites were located in Changins (46°19′ N 6°14′ E, 455m a.s.l), Delley (46°55′ N 6°58′ E, 494m a.s.l) and Utzenstorf (47°97′ N 7°33′ E, 483m a.s.l.). Climatic conditions and soil properties for the three sites and three seasons are described in Fig. S1-S3 and Table S1.

### Wheat varieties

We used 8 Swiss varieties which covered all quality classes: Molinera, Bodeli and CH 211.14074 with very high quality, Schilthorn, Falotta, Campanile and CH 111.16373 with high quality, and Colmetta with medium quality. All varieties were winter wheat, except for 211.14074 which is issued from the summer wheat breeding program, and Campanile which is a summer wheat but issued from the winter wheat breeding program. We chose varieties from the current Swiss national breeding program that were in use by farmers at the beginning of the project (i.e. 2020) or that were being developed, in order to have direct applicable results. Varieties were chosen based on their disease resistance profiles, as well as their morphological and agronomic characteristics to include differences in yield, protein content, foliage shape and awnness, with no more than 15 cm difference in height and no more than 5 days difference in phenological development to ensure synchrony in maturity. More detailed description of the agronomic characteristics of the varieties are provided in Table S2.

### Experimental communities

Experimental communities consisted of single variety plots, 2-varieties mixtures, and one plot with the 8 varieties mixed. We sowed every possible combination of 2-varieties mixtures, amounting to a total of 28 2-varieties mixtures treatments, to which we added the 8-varieties mixture. Each community was grown in a plot of 7.1 m^2^ (1.5m*4.7m). We replicated the experiment three times per site with the exact same variety and mixture composition. We used a complete randomized block design, with plots being randomized at each site within each block. Density of sowing was 350 viable seeds/m^2^. For the mixtures, seeds were mixed beforehand at a 2×50% ratio for 2-varieties mixtures and 8×12.5% for the 8-variety mixture in proportion to their weight. Plots were sowed mechanically each autumn (see Table S1) and fertilized with ammonium nitrate at a rate of 140 N/ha in 3 applications (40 at tillering stage – 60 at the beginning of stem elongation – 40 at the flag leaf stage). The trials were grown according to the Swiss *Extenso* scheme, i.e. without any fungicide, pesticide, and growth regulator (Finger & El Benni, 2013).

### Data collection

#### Phenology and height

For each plot, we recorded the heading date as the day of the year, in which 50% of the ears of the plot had fully emerged from the flag leaf. Plant height was measured in each plot at BBCH 59–75, by taking the average height in centimeters from the ground to the top of five random ears, excluding awns.

#### Light interception

In Changins (1260), light interception by the canopy was assessed with measurements of Leaf Area Index (LAI) with a ceptometer (Accupar LP-80, METER Group) three times during each growing season. We took the measures once early in the season (between days of year 117 and 120), once in the mid-season (days of year 137-146), and once at the end of the season before senescence (days of year 153-155). In each plot, three measurements were taken in the late morning by placing the sensor on the soil surface in three in-between rows (avoiding the in-between rows at the plot borders). Light measurements beneath the canopy were compared to ambient radiation through simultaneous PAR measurements of a calibration sensor, which was placed on a vertical post at 1.5 m above ground.

#### Traits measurements

At flowering time, we randomly sampled 6 healthy leaves per plot. We immediately wrapped this leaf in moist cotton; this was stored overnight at room temperature in open plastic bags. The following day, we removed excess surface water on the leaf and weighted it to obtain its water saturated weight (Cornelissen et al., 2003). This leaf was then scanned with a flatbed scanner (Perfection V39II, Epson), oven-dried in a paper envelope at 80°C for 72 hours, and subsequently weighed again to obtain its dry weight. Leaf Dry Matter Content (LDMC) was calculated as the ratio of leaf dry mass (g) to water saturated leaf mass (g). Using the leaf scans, we measured leaf area with the image processing software ImageJ (Schneider et al., 2012). Specific Leaf Area (SLA) was calculated as the ratio of leaf area (cm2) to leaf dry mass (g).

#### Ear density

Before harvest, we manually harvested horizontal bands of 1.5×0.3 square meters per plot. The location of the band was randomly chosen but we avoided plot edges (i.e. the band was located at more than 0.5m from the lower and upper edge of each plot). We counted the heads, and obtained ear density from the head counts.

#### Plot yield

At maturity, we harvested each plot with a combine harvester (Zürn 150, Schontal-Westernhausen, Switzerland). The harvested grains were dried when needed, weighed a first time, then sorted and cleaned by air and with a sieve cleaner, and subsequently weighted again. We measured hectoliter weight (test weight, HLW, kg/hl) and water content at the plot level using a Dickey-John machine (GAC 2100). Grain yield was subsequently standardized to 15% of humidity. Protein content (% of dry matter) was measured at the site level with a near-infrared instrument (ProxiMate^TM^, Büchi instruments). Thousand kernel weight (TKW, g) was measured at the plot level with a Marvin seed analyzer (GTA Sensorik, Neubrandenburg, Germany).

### Data analyses

#### Performance

Agronomic performance of the treatments was assessed with 5 response variables, namely grain yield (dt/ha), protein content (% of dry matter), Thousand Kernel Weight (TKW, g), Hectoliter Weight (HLW, kg/hl), and Zeleny sedimentation value (ml). For each of these variables and also for Leaf Area Index (LAI) and ear density, we calculated the corresponding overperformance in mixtures as the difference between observed and expected value of the mixtures, where expected value is the sum of the values in pure stands weighted by the relative abundance of each component (Loreau & Hector, 2001):

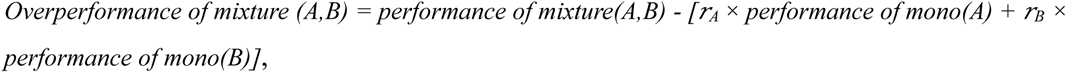

where *r_i_* indicates the relative abundance of the variety *i* in the mixture. The relative abundances were calculated based on the 50:50 mass ratio corrected with the thousand kernel weight values and germination rates of the varieties at sowing. This calculation was also performed for early and late Leaf Area Index (in Changins only).

The effects of year, site, and variety number on performance and overperformance were investigated using linear mixed-effects models with year interacted with site interacted with variety number nested into monoculture vs. mixtures as fixed factors, and replication per environments as well as variety composition as random factors, for instance:

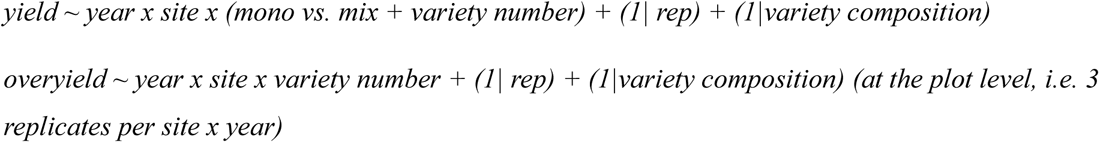

#### Stability

We quantified the stability of each response variable using Weighted Average of Absolute Scores from the BLUP decomposition (WAASB) developed by Olivoto (2019).This was chosen as BLUP has been shown to be the most predictively accurate model, and the WAASB index presents several advantages and increased robustness compared to AMMI-based stability indexes (Olivoto, Lúcio, da Silva, Marchioro, et al., 2019).

The effects of year/site and variety number on stability were investigated with a similar linear mixed-effects models, with year or site (depending on temporal vs. spatial stability) interacted with variety number (2 or 8) nested into monoculture vs. mixture as fixed factors, and variety composition as random factors, e.g.

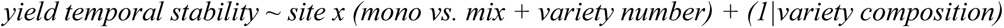

For overall stability, we simply used a linear model with mono vs. mix and variety number as factors.

#### Multi Traits Performance and Stability

Multitrait performance and stability was assessed using the Multi Trait Stability Index developed by Olivoto (2019). The index was computed using the five response variables described above, with equal weights for each: grain yield, protein content, TKW, HLW, and Zeleny sedimentation value. Performance and stability were given equal weights in the index calculation. The genotype/mixture with the lowest MTSI is the closest to the ideotype and presents a high performance and stability for all analysed variables (Olivoto, Lúcio, da Silva, Sari, et al., 2019).

#### Monoculture characteristics

for each 2-variety mixture, we calculated the absolute and relative difference in yield, protein content, height, heading date, density, Leaf Area Index (in Changins only), SLA (in Changins only) and LDMC (in Changins only) between the two components when grown in pure stands as:

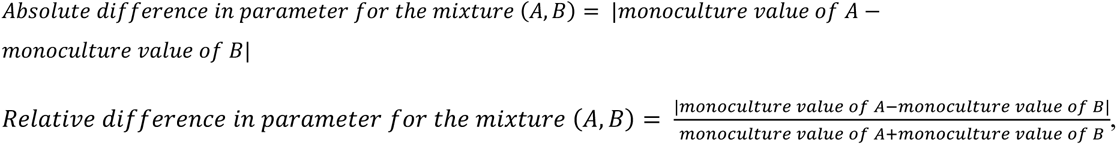

These indices allow to quantify how different the varieties in the mixtures are (Stefan et al., 2024). The indices were computed per site and year, and subsequently averaged across sites and/or years when needed. The absolute difference was used in models investigating absolute response variables (such as overyielding, for instance), while the relative difference was used in the stability models, as the relative difference allows to compare between environments more easily.

To investigate relationships between overperformance and explanatory parameters, we used linear mixed-effects models with absolute differences in yield, protein content, height, heading date, density, LAI, FPAR, SLA and LDMC (in Changins only), as well as awns difference (yes when one variety has awns, the other not; no when both varieties have awns or both do not) as fixed factors, and site within year, variety composition as random factor, e.g.

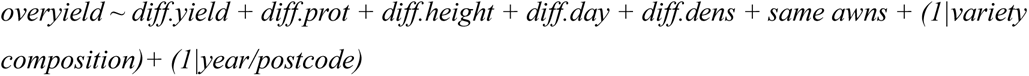

We used this model as we were interested to see what characteristics were linked to overperformance across years and sites, i.e. that were not environment-dependent. This model was run for 2-varieties mixtures only.

For stability responses, we used the same type of model but averaged the relative differences to the scale of stability investigated (spatial, temporal, overall).

Homogeneity of variance and normality of residues for linear mixed models were assessed visually and with Shapiro–Wilk tests (Royston, 1982). Effect sizes were calculated from marginal means obtained using the function *emmeans*, and post-hoc pairwise comparisons were computed using Tukey test from the *emmeans* function (Lenth, 2021). Effect sizes are indicated in brackets in the text.

## Results

### Performance and overperformance

In a first step, we looked at how the response variables were affected by environmental and experimental factors (i.e. year, site, and diversity treatment). Grain yield was significantly affected by year*site, and year*site*variety number (Table S3, Fig. S4, Fig. S5). Post-hoc comparisons showed however that there was no significant difference in yield due to variety number (i.e. 2 ou 8 varieties) within any year*site. Protein content was only affected by year and site, just as TKW and HLW (Table S3). Zeleny sedimentation value significantly responded to site and monoculture vs. mixtures: across all varieties and combinations, Zeleny sedimentation value increased in mixtures compared to monocultures (from 51.5 to 52.4 mL, F-value = 5.19, p-value = 0.023, Table S3, Fig. S6).

The same models were applied to overperformances (i.e. the difference between observed value in the mixture and the expected value based on the corresponding pure stands) and showed that for all parameters, overperformance values were affected by year and site, but not by variety number (Table S4, Fig. S7, Fig. S8). Overyield and overprotein content were affected by year*site, while overTKW only by year and overHLW only by site. OverZeleny was greatly impacted by year and site, with lowest values in 2021 (−0.5 mL in 2021 vs. 2.7 mL in 2022) and highest in Utzenstorf (3.15 mL). In addition, overZeleny had a significant site*variety number factor, but posthoc comparisons did not show any effects of variety number within site.

When looking at the average overperformance deviation from 0 (Table 1), we found that overprotein was significantly negative across all sites and years (−0.11% in average, Table 1), while overZeleny was significantly positive (+1.08 mL in average, Table 1). OverZeleny was particularly positive in Delley and Utzstenstorf, as well as in 2022 and 2023 (Table 1), but negative in Changins 2021 (Table S5). Overprotein was significantly negative in Delley and in 2021; the only significant positive value was observed in 2022 in Changins (Table S5). Results for overyield, overTKW and overHLW were more contrasted, with notable positive overyield in Changins and in 2021, but negative in Delley 2022 and Utzenstorf 2022. OverTKW was globally negative in 2021 and positive in 2022, while overHLW was positive in Delley 2022 and Utzenstorf 2022 but negative in Changins 2022 and Utzenstorf 2023. Early overLAI were never significantly positive or negative, but late overLAI was significantly positive in 2021, 2023, and in the global average. Overdensity was significantly positive in Changins and negative in the other two sites.

**Table 1:** Results of the t-test to evaluate whether overperformance is significantly different from 0. Numbers indicate the average absolute values per site (Changins, Delley, Utzenstorf) and per harvest year (2021, 2022, 2023). Bold numbers indicate environments where the t-test was significant.

|  | *Changins* | *Delley* | *Utzenstorf* | *2021* | *2022* | *2023* | *Average* |
| --- | --- | --- | --- | --- | --- | --- | --- |
| *Overyield* | **1.7** | -0.9 | -0.5 | **0.59** | -0.34 | -0.01 | 0.08 |
| *Overprotein* | 0.023 | **-0.3** | -0.06 | **-0.17** | -0.07 | -0.1 | **-0.11** |
| *OverTKW* | 0.07 | 0.07 | -0.056 | **-0.29** | **0.22** | 0.16 | 0.029 |
| *OverHLW* | -0.55 | 0.016 | 0.095 | 0.078 | -0.066 | -0.44 | -0.14 |
| *OverZeleny* | -0.15 | **1.5** | **1.86** | 0.035 | **2.18** | **1.03** | **1.08** |
| *OverLAI early* | 0 | NA | NA | 0.03 | 0.14 | -0.07 | 0 |
| *OverLAI late* | **0.3** | NA | NA | **0.55** | 0.035 | **0.14** | **0.3** |
| *Overdensity* | **21.7** | **-12.9** | **-11** | -11.1 | 7.9 | 0.55 | 0.75 |

In a second step, we looked at relationships between overperformances and possible explanatory variables, namely differences in monoculture characteristics and differences in awns. Results are detailed in Table S6, S7 and S8, summarized in Table 2, and show that a few general patterns emerge.

**Table 2:** Effects of explanatory variables on overperformance parameters. Filled boxes indicate a significant positive (+) or negative (-) effect based on the linear mixed-effects model.

|  | *Overyield* | *Overprotein* | *OverTKW* | *OverHLW* | *OverZeleny* |
| --- | --- | --- | --- | --- | --- |
| *Awns difference* |  |  |  |  |  |
| *Diff in mono yield* |  | **+** |  |  | **+** |
| *Diff in mono protein* |  | **-** |  |  |  |
| *Diff in mono height* | **-** |  |  | **-** |  |
| *Diff in mono heading day* |  |  | **-** |  | **-** |
| *Diff in mono density* |  | **+** |  |  |  |
| *Diff in mono LAI early* |  |  |  |  |  |
| *Diff in mono LAI late* |  |  |  |  |  |
| *OverLAI early* |  | **+** |  |  | **+** |
| *OverLAI late* | **+** |  | **-** |  |  |
| *Diff in mono SLA* |  |  |  |  |  |
| *Diff in mono LDMC* |  | **-** |  |  |  |

First, monoculture difference in height had a significant effect on overyield and overHLW (Table S6). This effect was negative for both overyield (estimate = −0.26, Fig.1) and overHLW (−0.07), indicating that mixture benefits for yield and HLW were higher when the two varieties mixed did not have a large difference in height when grown in monoculture.

**Fig. 1:**
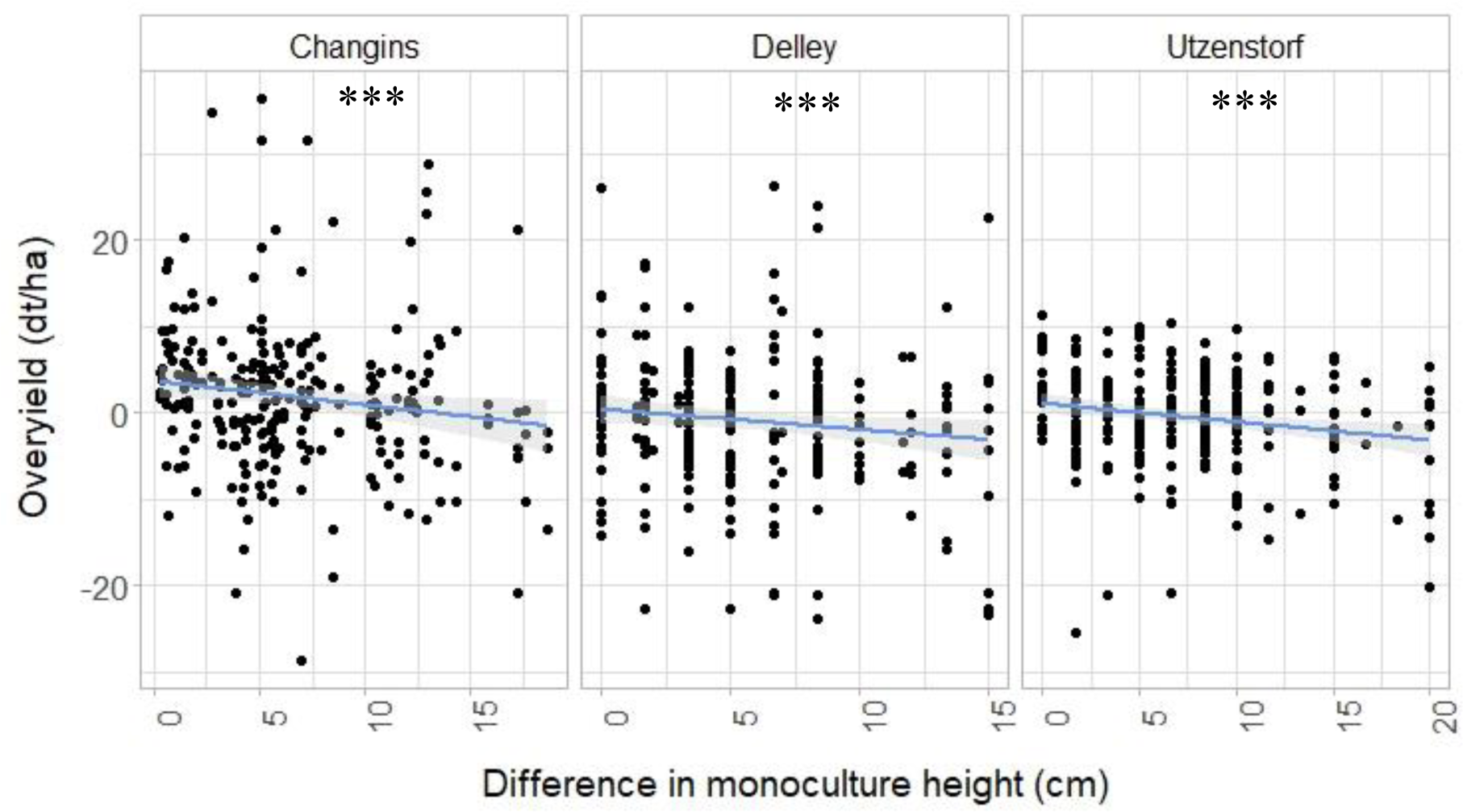
Grain overyield (dt/ha) of the mixtures in relationship to the mean difference in height of the corresponding varieties when grown in monocultures (cm), in Changins, Delley, and Utzenstorf. n=754. The lines represent linear regression fittings, with the grey area representing the 0.95 confidence interval. Stars represent significant relationships at p-value < 0.05.

Secondly, monoculture difference in heading day significantly affected overTKW (−0.07) and overZeleny (−0.19) (Table S6). In the two cases, the relationship was negative, indicating that benefits were higher when mixing varieties with similar phenologies and heading dates.

Thirdly, monoculture difference in yield had a positive effect on overprotein (0.015) and overZeleny (0.1) (Table S6), suggesting that mixture benefits for protein content and Zeleny sedimentation value were higher when mixing varieties with greater differences in monoculture yield.

The remaining effects were less clear and varied across response variables; for instance, difference in protein content between the mixed varieties had a significant negative impact on overprotein (−0.11), indicating that a lower difference in protein content between the two varieties correlated with increased overprotein. We also observed a positive relationship between overprotein and monoculture difference in ear density (0.002), indicating a larger mixture benefit for protein content when mixing varieties with larger differences in ear density when grown as pure stand. Mixing varieties with different awning characteristics did not affect any of the overperformance response variables.

The additional measures taken in Changins allowed to gain more insights into the role of light for all the response variables except HLW (Table S7). Specifically, the response variables showed strong relationships with either early or late overLAI; notably overprotein (0.46) and overZeleny (1.15) positively correlated with early overLAI, while overyield (1.9) positively correlated with late overLAI (Fig. 2). This indicates that overperformance was generally higher in mixtures with a higher overLAI, i.e. in mixtures where the LAI was larger than the relative sum of the corresponding LAIs in pure stands. There was one exception to this general result for TKW, with overTKW negatively correlating with late overLAI (−0.36). Finally, difference in LDMC negatively affected overprotein (−9.3), suggesting increased mixture benefits when combining varieties with similar LDMC.

**Fig. 2:**
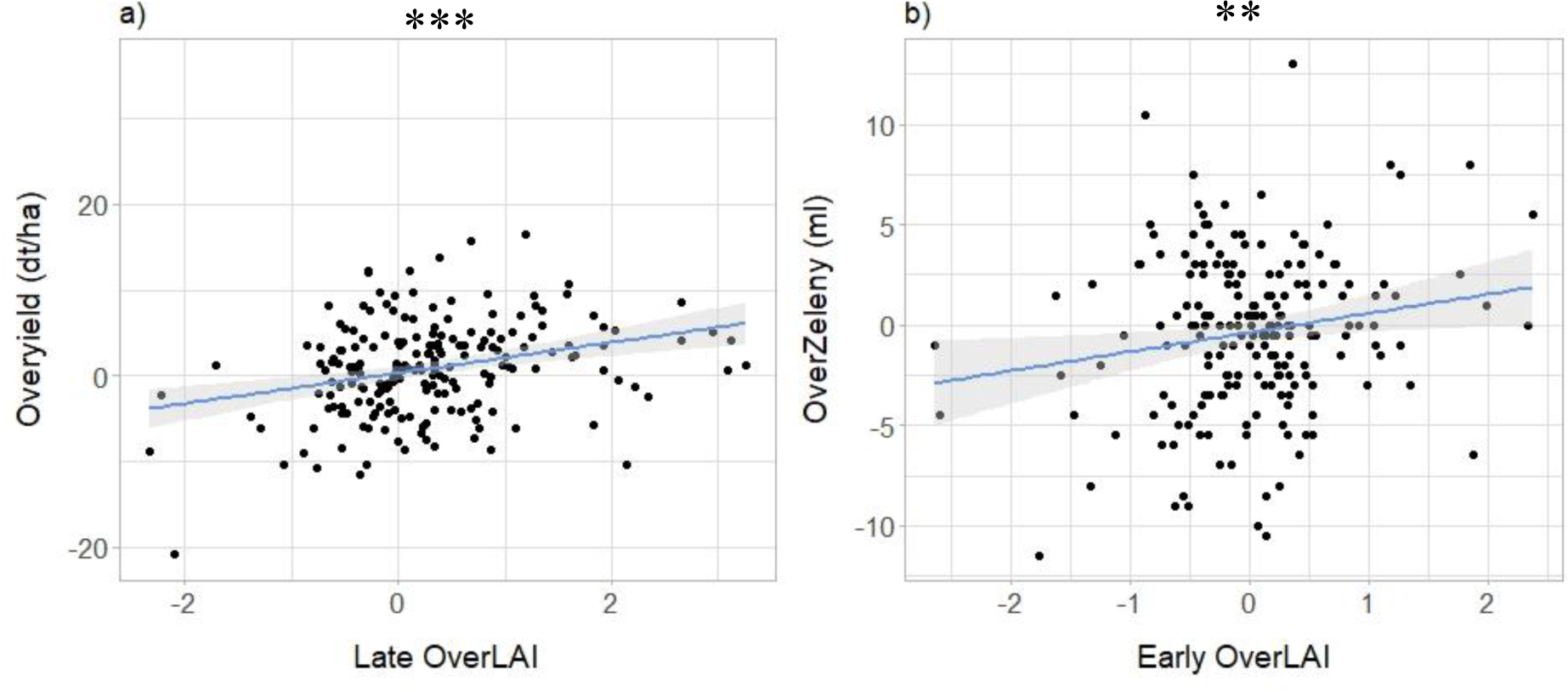
Grain overyield (dt/ha) (a) and overZeleny sedimentation value (mL) (b) of the mixtures in relationship to overLAI (Leaf Area Index) in Changins. n=246. The lines represent linear regression fittings, with the grey area representing the 0.95 confidence interval. Stars represent significant relationships at p-value < 0.05.

To summarise, Table 2 allows to show that overprotein was the most responsive to our explanatory variables, followed by overZeleny. Overyield and overTKW were less affected by our explanatory variables, while overHLW only responded to one criterium.

### Temporal stability

Stability per site was assessed with the score WAASB for each parameter, calculated per site across the 3 study years. It thus represents temporal stability. A lower WAASB score indicates more stability.

Yield stability score was not affected by site or diversity treatment (Table S9). Protein stability score (WAASB protein) was significantly affected by site and monoculture vs. mixture: WAASB protein was lowest in Delley, followed by Utzenstorf (+62.5%) and Changins (+62.5%, Fig. S9), and significantly lower in mixtures compared to monocultures (−14%, Fig. 3a). For TKW as well, we found a significant increase in stability (i.e. a decrease in WAASB TKW) in mixtures compared to monocultures: the stability score went from 0.322 in monocultures to 0.209 in mixtures (−35%) (Fig. 3b). WAASB TKW was also significantly lower in Changins compared to Delley (−30%, Fig. S9).

**Fig. 3:**
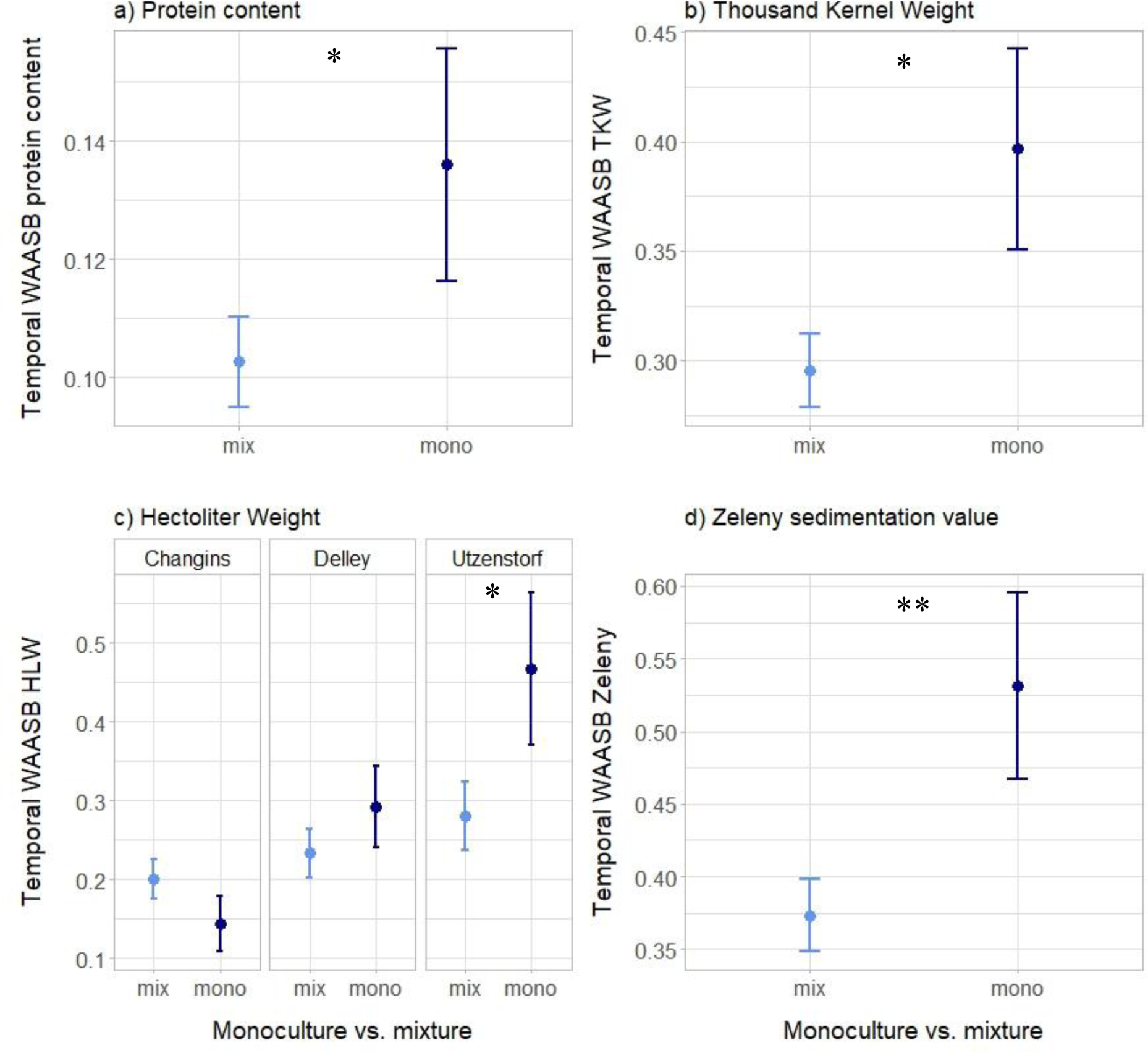
Temporal Stability scores for protein content (a), TKW (b), HLW (c), and Zeleny (d) in response to monoculture vs. mixture, and to site for HLW. n=111. Lower WAASB scores indicate higher stability. Dots represent the mean values across plots; lines represent the standard error. Stars placed above or next to the results represent the significance of monoculture vs. mixture.

Temporal stability for HLW was affected by site interacted with monoculture vs. mixture: specifically, we found that in Utzenstorf, WAASB HLW was 57% lower in mixtures compared to monocultures (Fig. 3c). Zeleny stability responded to site and monoculture vs. mixture: the stability score was significantly higher in Utzenstorf compared to Changins (−27%) and Delley (−35%, Fig. S9). It was also significantly lower in mixtures compared to monocultures (Fig. 3d), with a reduction of 21%, indicating an increase in temporal stability of Zeleny in mixtures.

When investigating into explanatory mechanisms, we did not find many significant effects (Table S10): there was only an effect of awns difference on WAASB protein, with a lower stability score (i.e. increased stability) when the two varieties combined had similar awn characteristics (−30% in comparison to mixtures with one variety awned and the other not). There was no effect of any additional light variables measured in Changins (Table S11).

### Spatial stability

Stability per year was assessed with the score WAASB for each parameter, calculated per year across the 3 study sites. It thus represents spatial stability. As before, a lower WAASB score indicates more stability.

Stability in yield was affected by year interacted with monoculture vs. mixture (Table S12): posthoc comparisons however did not show any effect of diversity within year. Stability in protein content was higher in 2021 than 2023 and 2022: WAASB protein was 87% lower in 2021 compared to 2022, and 81% lower in 2021 compared to 2023 (Fig. S10). Regarding thousand kernel weight, WAASB TKW was significantly lower in mixtures compared to monocultures (−41%, Fig. 4a), showing an increase in spatial stability of TKW in mixtures. Stability in HLW and Zeleny were both only affected by year: WAASB HLW was higher in 2023 than 2022 and 2021 (+16%), while WAASB Zeleny was significantly higher in 2022 compared to 2023 (+42%. Fig.S8). There was no effect of any of the explanatory variables on spatial stability (Table S13).

**Fig. 4:**
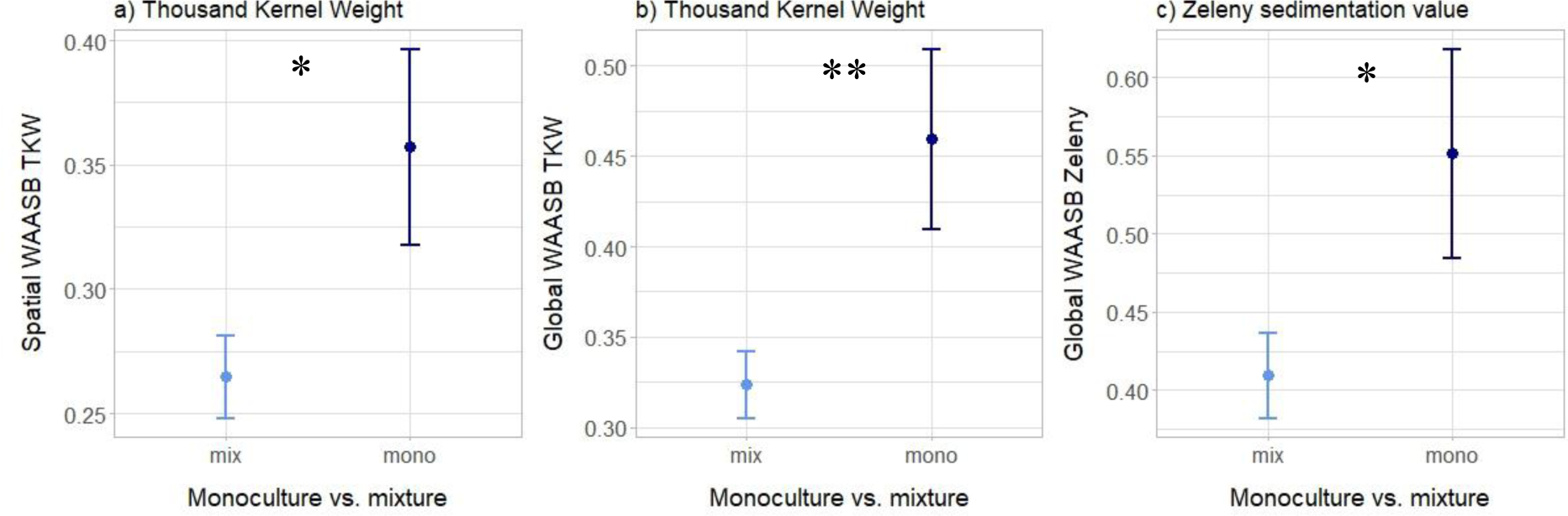
Spatial Stability score for TKW (a), Global Stability score for TKW (b) and Zeleny (c) in response to monoculture vs. mixture. n=111. (a), n=37 (b,c) Lower WAASB scores indicate higher stability. Dots represent the mean values across plots; lines represent the standard error. Stars placed above or next to the results represent the significance of monoculture vs. mixture.

### Global stability

Global stability was assessed with the score WAASB for each parameter, calculated across the 9 environments of the study (i.e. across sites and years). It thus represents overall stability. As before, a lower WAASB score indicates more stability.

In the case of thousand kernel weight and Zeleny sedimentation value, we found a reduction in WAASB score in mixtures compared to monocultures (−42% for WAASB TKW and −13% for WAASB Zeleny, Fig. 4b and 4c, Table S14), indicating an increase in stability for TKW and Zeleny in mixtures.

When looking at explanatory variables, we found that WAASB TKW and WAASB HLW were positively affected by difference in monoculture height (3.5 and 2.8, Fig. 5a and 5b, Table S15), indicating that mixtures were more stable for TKW and HLW when combining varieties with similar heights in monocultures. WAASB yield was negatively affected by difference in monoculture protein content (−1.8) (Table S15).

**Fig. 5:**
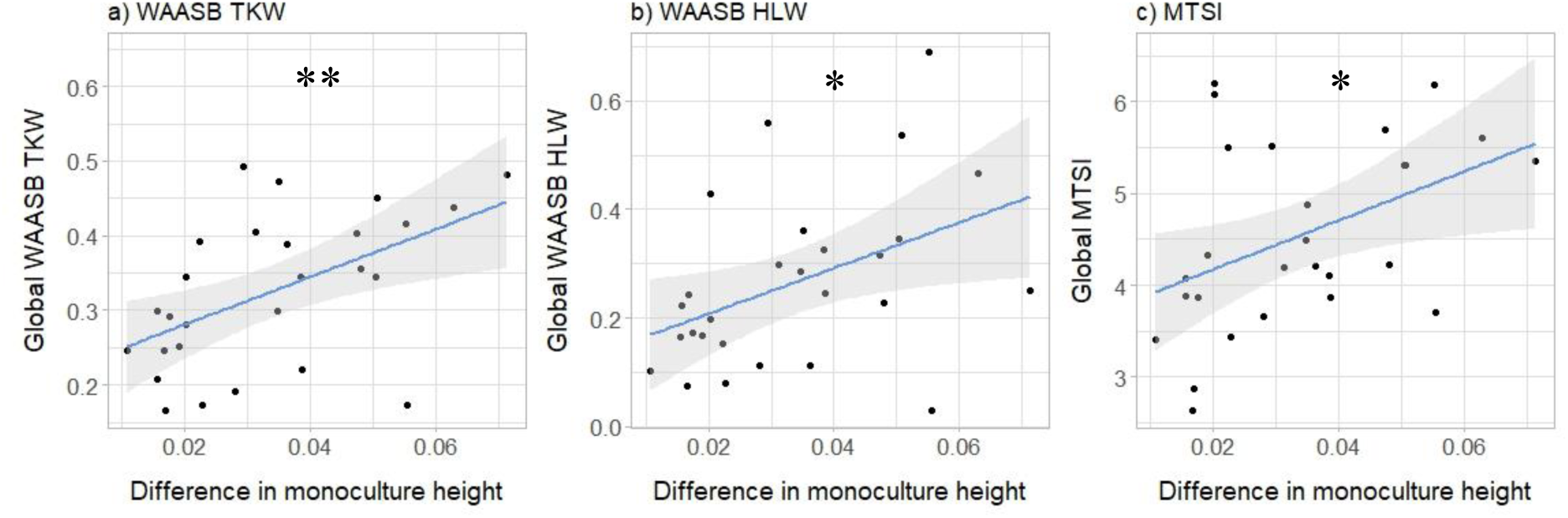
Global Stability scores for TKW (a) and HLW (b), as well as Global Multitrait Stability Index (c) of the mixtures in relationship to difference in monoculture height. n=28. Lower WAASB scores indicate higher stability. The lines represent linear regression fittings, with the grey area representing the 0.95 confidence interval. Stars represent significant relationships at p-value < 0.05.

### Multitrait performance and stability

MTSI is a combined measured of multitrait performance and stability. Lower MTSI indicates closer proximity to the ideal genotype, and thus, higher performance and stability.

Temporal MTSI (i.e. MTSI per site, calculated across years) was significantly affected by site and monoculture vs. mixture (Table S9). Notably, temporal MTSI was lower in Changins compared to Delley and Utzenstorf (−24%, Fig. S11) and lower in mixtures compared to monocultures (−11%, Fig. 6). This indicates that across years, mixtures were performing better and were more stable than monocultures. There was no effect of the explanatory variables (Table S10, S11).

**Fig. 6:**
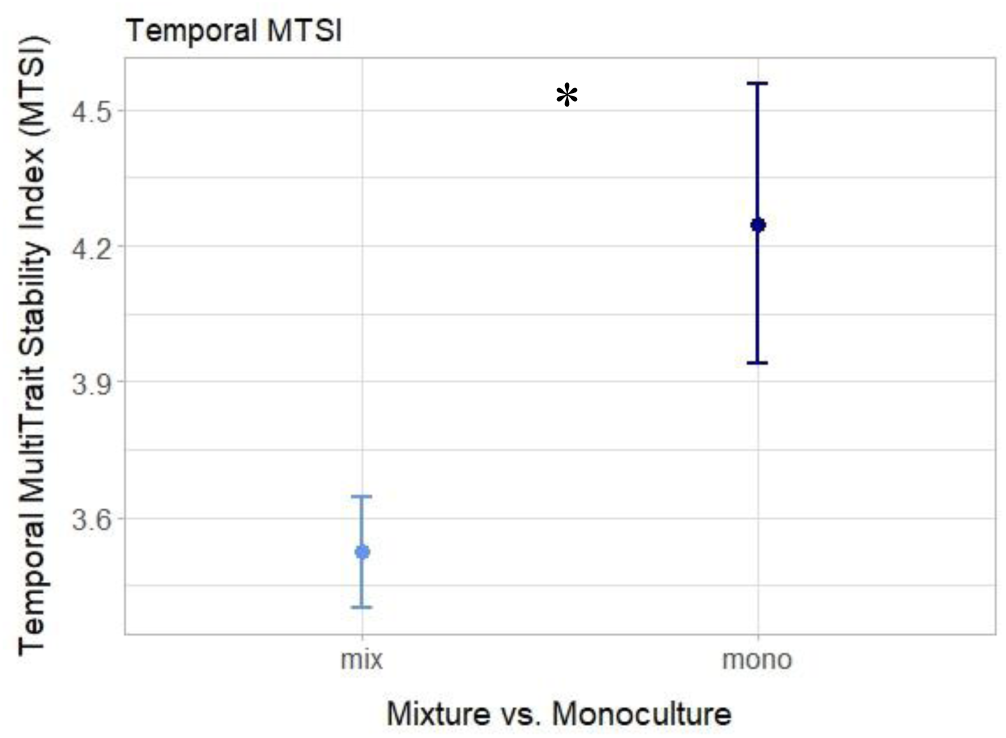
Temporal Multitrait Stability Index in response to monoculture vs. mixture. n=111. Lower MTSI scores indicate higher stability and performance. Dots represent the mean values across plots; lines represent the standard error. Stars placed above or next to the results represent the significance of monoculture vs. mixture.

Spatial MTSI (i.e. MTSI per year, calculated across sites) was affected by year and year interacted with number of varieties (Table S12): MTSI was lower in 2023 than 2021 (−25%) and 2022 (−17%, Fig. S11). There was no significant effect of variety number within any year, nor any effect of the explanatory variables (Table S13).

Global MTSI (calculated across sites and years) was positively affected by difference in monoculture yield (2.1, Fig. 5c) and difference in monoculture height (4.7, Fig. 5c) (Table S15), indicating that overall performance and stability was higher in mixtures combining varieties with similar heights and yields when growing in monocultures. All mixtures and monocultures were ranked according to their global MTSI score; this result can be visualized in Figure 7.

**Fig. 7:**
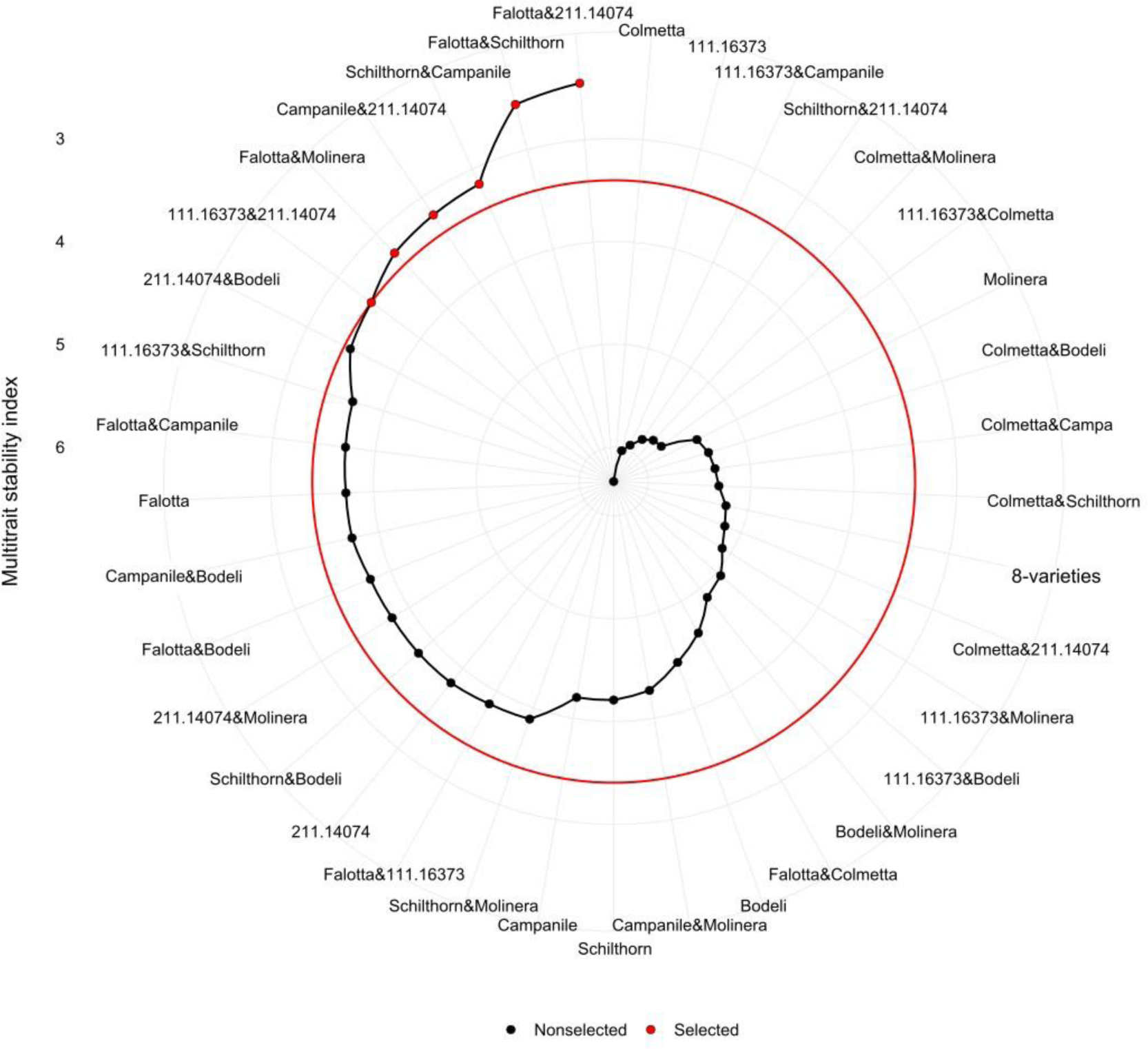
MTSI ranking of all mixtures and monocultures. Lower MTSI scores indicate higher stability and performance. The red line represents the 15% selection intensity (i.e. red dots represent the 15% best genotypes/mixtures).

## Discussion

In this study, we used a multitrait approach to investigate the effects of variety mixtures on performance and stability of several parameters reflecting crop quantity and quality in wheat production systems. We expected higher performance and stability of these parameters in mixtures compared to monocultures. We found evidence in this direction for Zeleny values and the stability of several parameters; in addition, when combining the stability and performance of all parameters, we showed that mixtures outperformed monocultures. In a subsequent step, we investigated links between performance and stability of these parameters and traits of the mixture components. While many relationships depended on environmental context, we did find that in general, the best mixtures among our samples were those combining varieties with similar heights and phenologies, but different yields when grown in pure stands. In addition, mixture performance was higher when the combined varieties were able to intercept more light than in monocultures.

### Mixtures generally outperform monocultures

In order to have a comprehensive view of both crop quantity (i.e. yield) and crop agronomic quality, we considered five crop parameters in this study, namely grain yield, grain protein content, Thousand Kernel Wight (TKW), Hectoliter Weight (HLW), and Zeleny sedimentation value. These were chosen because they are important to farmers and wheat processors, and are part of standard evaluations by processors, industry, and the whole wheat value chain (Kleijer, 2002; Saurer et al., 1991; swissgranum, 2023b). When considering all the response parameters, our study showed that mixtures outperformed monocultures in terms of overall performance and stability. Notably, the temporal multitrait stability index – which combines all parameters and evaluates both performance and stability – was lower in mixtures than in monocultures (Fig. 6), indicating that at each site, mixtures performed better and/or were more stable across years than monocultures. Spatial MTSI and global MTSI also showed similar trends, but these were not significant (see Fig. S12), probably due to small sample sizes for global MTSI (n=37). We could nonetheless rank all mixtures and monocultures according to their global MTSI scores and provide several options for promising mixtures to farmers (Fig. 7). This multitrait result is consistent with Lazzaro et al. (2018), who showed that overall crop performance of mixtures was higher than individual varieties when considering several agroecosystem services together. Similarly, Hoang et al. (2021) demonstrated that mixtures have the potential to improve overall crop performance (including grain quality). This result is all the more staggering considering that our study was a full diallel design, meaning that all the 2-variety combinations were considered: we did not specifically focus on chosen, smartly designed mixtures. This indicates that even when mixing varieties randomly, chances are that the mixtures could bring benefits.

In our case, lower MTSIs in mixtures were largely driven by lower WAASB scores of several response parameters: this was notably the case for the stability scores of protein content, TKW, and Zeleny (Fig. 3, Fig. 4), highlighting the benefits of variety mixtures for increased stability in these parameters. Similar results were found for instance by Tremmel-Bede (2016), who showed that variety mixtures and composite cross populations have a stabilizing effect on wheat quality parameters, or Döring (2015), who demonstrated that mixtures had more stable protein yields than pure lines (Döring et al., 2015).

In addition to benefits for stability, our study also showed that mixtures outperformed monocultures for Zeleny sedimentation value. This was the case for absolute values, i.e. mixtures had a higher Zeleny value than monocultures across sites and years (Fig. S6), but also for relative terms, i.e. overZeleny was significantly positive overall (Table 1). This means that in mixtures, the Zeleny value was generally higher than the relative sum of the corresponding monoculture values. This finding is rather new, as very few studies have looked at Zeleny sedimentation values in mixtures vs. monocultures, and in the few that did, no difference in Zeleny was found (Hoang et al., 2021; Osman, 2006). Zeleny is an important characteristic for bread baking and is standardly used, among other parameters, by breeders and processors to select high baking quality lines (Escarnot et al., 2018) and predict final behaviour during the baking process (Hildermann et al., 2009). Higher Zeleny values are generally associated with both higher gluten content and a better gluten quality, which strongly influences final bread volume and quality (Hrušková & Faměra, 2003). Therefore, higher Zeleny values in mixtures could potentially lead to increased baking quality, which could be of interest for wheat breeders, processors, bakers, and consumers.

### But no one-size-fits-all solution: the success of mixtures is dependent on the assessed parameters

While mixtures outperformed monocultures when combining all the assessed parameters, this was not always the case when looking at each parameter individually. For instance, in our experiment, overZeleny was positive, overyield, overTKW and overHLW were not significantly positive nor negative, while overprotein content was even negative across all environments. This study therefore emphasizes the importance of considering several response parameters to properly assess mixture performance.

While contrary to our initial hypotheses, the absence of positive overyielding is still consistent with the literature, where several studies found that yield advantages often vary according to the mixture or the environmental conditions (Alsabbagh et al., 2022; L. Li et al., 2023; Mengistu et al., 2010). Li (2023) did for instance show that increasing the number of varieties does not always promote productivity, which is in line with our results of yield (Table S3 and S4). Alsabbagh (2022) and Hoang (2021) even found that increasing genetic diversity had a generally negative effect on yield, while Mengistu (2010) only found small yield advantages in mixtures that were rarely significant. Importantly, while we did not find significant yield benefits, we also did not find yield losses in mixtures, which suggests that mixtures do not penalize farmers in terms of grain yield (Creissen et al., 2016).

Mixtures had an overall negative effect on protein content, which was unexpected: in Hoang (2021) for instance, variety mixtures improved protein content. This negative result for protein content is all the more surprising when considering that mixtures had an overall positive effect on Zeleny, and that these two parameters are often positively correlated (Pasha et al., 2007). A positive effect on Zeleny combined with a negative effect on protein content thus shows that the benefits for Zeleny are independent from protein content, and suggest a true increase in protein quality in wheat mixtures. This finding is rather new; further research should be conducted to confirm this result, and baking tests should be performed to check whether this increase in quality holds true up until the actual bread baking.

Stability results were more consistent when looking at each parameter individually, with increased stability in mixtures for protein, TKW, HLW, and Zeleny. Among the few studies looking at stability of crop quality in mixtures, there were similar findings: Tremmel-Bede (2016) observed positive effects on the stability of quality traits in mixtures, while diversity was beneficial for grain weight in Alsabbagh (2022). Surprisingly, we did not find significant mixture benefits for yield stability in our study, which was contrary to our expectations based on previous literature from both ecological and agronomical fields (Huang et al., 2024; Kaut et al., 2009; Mengistu et al., 2010; St. Luce et al., 2020; Tilman & Downing, 1994; Tremmel-Bede et al., 2016). This suggests that it is not just diversity *per se* that matters, but also mixture composition and environmental conditions (Schöb et al., 2015; Tilman et al., 1997): if there is no abiotic or biotic stressors, or if the variety responses to these stressors are not different enough, then the potential compensatory effects in the mixtures might not take place (Stefan et al., 2024).

### Explaining performance and stability of mixtures from variety traits

One of our main hypotheses was that mixing varieties that are more different – in terms of agronomical traits (yield, protein content, density) but also morphological (height, awns), phenological (heading date) or functional (LAI, SLA, LDMC) traits – would lead to mixtures with increased benefits. This was not necessarily true in our study: we observed as many positive effects of variety differences on overperformance than negative effects (see Table 2). When looking into more details, we had rather contrasted results depending on the variety trait investigated or the response parameter of interest. This shows once again that there is no simple, universal solution for increased performance of all parameters in diverse crop communities (Brooker et al., 2015). Nevertheless, a few important points emerged among our results. First, this study underlined the importance of height as a key variety trait, with lower differences in height between varieties correlating with better overall mixture performance and stability (MSTI, Fig. 5c), increased benefits for yield and HLW (Table 2), as well as for stability of TKW and HLW (Fig. 5a and 5b). Mixing varieties that had similar heights when grown in monocultures thus led to higher benefits. This result is confirming previous research such as Gawinowski (2024), in which the mixtures with varieties more different in height also underyielded. Height is a trait for which plants generally demonstrate adaptive similarity: in response to competition for light and to avoid shade, all individuals tend to converge to the same height in a crop field (Dahlin et al., 2020; Stefan et al., 2022). Smaller varieties thus need to compensate and generally over-invest in vegetative organs at the expense of reproductive ones, thereby reducing their yield potential (Anten & Vermeulen, 2016; Wille et al., 2017).

The observed significant role of height points us towards the crucial role of light interception for improved crop performance (Gawinowski et al., 2024). The importance of light was confirmed by the predominant role of LAI in our mixtures (Table 1 and 2): late overLAI was significantly positive in mixtures, indicating that mixtures intercepted more light than what was expected based on their respective components in pure stands. This is consistent with many studies looking at diverse agricultural systems, notably with Huang et al. (2024), who found that cultivar mixtures generally increased leaf area index in various crop systems. Engbersen (2022b, 2022a) and Board (1992) also observed greater light interception in species mixtures, and identified this as a driving mechanism of yield benefits. As expected, in our case, increased overLAI correlated with increased benefits for yield in variety mixtures (Table 2). This was also the case in Engbersen (2022) for species mixtures, and in Hu (2019) for maize, where mixed cropping optimized canopy structure and promoted higher grain yield. Interestingly, overLAI in mixtures did not only positively correlate with overyield, but also with increased benefits for Zeleny and for protein. This shows that higher light interception does not only promote higher grain yield, but also improves grain quality. This statement is further supported by our results of ear density, which showed that mixing varieties with more different densities was more beneficial for protein (Table 2). Ear density is closely related to tillering ability and therefore also reflects the potential of varieties to modulate their light interception in response to light availability and other environmental constraints (Gawinowski et al., 2024; Maddonni et al., 2001; Weiner et al., 2010). For instance, plasticity in ear numbers and tillering was found as the main contributor to variation in mixture yields in the study of Gawinowski. In our study, we could unfortunately not assess trait plasticity or monitor tillering and ear numbers of each individual variety within the mixtures, as it was impossible to distinguish every component from each other.

Secondly, our study emphasized the importance of closer phenologies for increased performance of the mixtures. This was unexpected, as it has been largely suggested in the literature that some benefits of variety mixtures stem from phenological buffering, i.e. mixtures buffer the risk of flowering “too early or too late” (Fletcher et al., 2019). While large differences in phenology are not desirable for technical and harvest considerations, smaller phenological differences can in theory be beneficial to minimize risks of extreme weather events happening at a critical development stage (e.g. heat or frost event during flowering) (Haghshenas et al., 2021; Tschurr et al., 2023). We did not find any support for this theory in our case; this might be because there was no particular stress during critical development stages, or because the mechanisms of phenological buffering are more complicated than what current theory suggests. Similar findings were reported by Haghshenas (2021) in a situation of post-anthesis water stress: under stressful conditions, they even reported lower heterogeneity in the ripening pattern of mixtures.

Finally, we found improved quality in terms of protein and Zeleny when mixing varieties with larger differences in yield. Thus, mixing a high-yielding variety with a low-yielding variety can increase quality benefits. One explanation to this is that high-yielding varieties tend to have lower quality than low-yielding varieties – due to trade-offs in resource allocation and breeding histories (Anderson et al., 1998; Michel et al., 2019; Simmonds, 1995) – and combining the two can therefore mask the lower quality of the high-yielding variety and boost overall quality (Zhou et al., 2014).

We only assessed aboveground traits in this study, which are substantially linked to light interception, and this can be why we mostly observed a significant role of light interception mechanisms. Belowground processes, such as root complementarity (Freschet et al., 2021; Stomph et al., 2020), plant-microbiome interactions (Duchene et al., 2017; Stefan, Hartmann, et al., 2021), or plant-soil feedbacks (Eroğlu et al., 2024) can also play a role in driving complementarity and productivity. In addition, belowground processes can provide better insights into competition for plant nutriments or water (Engbersen et al., 2021; Foxx & Fort, 2019; Manoli et al., 2017; Song et al., 2009), which can then influence crop growth, grain yield and grain quality. In our experiment, we could not well explain variations in overTKW and overHLW, and maybe this is due to the importance of water availability – which we did not measure – for grain filling (Royo et al., 2007; Torrion & Stougaard, 2017). Finally, the bulk of our measures were done only once per season and do not account for temporal variation in growth or resource partitioning. Only LAI was measured several times in the season, and results show that there are indeed changes between the early measures and the late measures: for instance, in the early season, overLAI correlated with overprotein content, while in the late season, it correlated with overyielding. This shows that different plant processes happen at different timeframes of the growing season (Trinder et al., 2012; Zhang et al., 2017); the moment of peak growth rate might not be the same as peak grain filling, and some resources might be more important during vegetative growth while others might matter more during grain formation (Ashraf & Bashir, 2003). This temporal differentiation of processes and resource use might thus be crucial to explain overperformance (Engbersen et al., 2021; Zhang et al., 2015).

### Effects on environmental conditions on mixtures stability and performances

The results described above are valid across environments in our study; they represent the general effects and links that we could find among the great variability partly due to natural heterogeneity inherent to field trials, but mostly due to changing environmental conditions. Indeed, performance and stability of our wheat communities were generally strongly dependent on environmental conditions, i.e. year and site. This was expected based on previous literature (e.g. see (Döring et al., 2015; Su et al., 2023; Weih et al., 2021)). Notably, we expected increased performances and stability of mixtures in more stressful environments, following the stress-gradient hypothesis (Bertness & Callaway, 1994; Lortie & Callaway, 2006). When looking at all the response parameters together, we observed lower MTSI scores in Changins and in 2023, indicating a better overall performance and stability in these environments. Overyielding was also positive in Changins only (Fig. S7), while TKW and HLW were more stable there (Fig. S9). Environmental conditions were slightly harsher in Changins compared to the two other sites: the soils in Changins were characterized by higher than optimal pH for wheat cultivation (pH > 7) (Mahler & McDole, 1987; Vitosh, 1998), while precipitations during early season were more elevated (Fig. S1). On the contrary, in Utzenstorf, the most productive environment (Fig. S4, highest yielding site), none of the response parameters were very stable. These results may hint us towards higher yield benefits and stability in lower productive conditions.

2023 was marked by very hard conditions regarding drought and heat stress (Fig. S1-S3) and was the lowest yielding year (Fig. S4). Yield and Zeleny were particularly more stable in 2023 (Fig. S10), but this was not the case for all parameters: protein for instance was more stable in 2021. Moreover, regarding overperformance, nothing was positive in 2023 except for overZeleny (Table 1). There was no significant overyielding in 2023, even though this was the most stressful year in terms of productivity; this illustrates the complexity of environmental effects. Similar results were previously reported, with Alsabbagh finding little support for the stress gradient hypothesis (Alsabbagh et al., 2022), or Döring who observed that yield benefits were inconsistent across years and sites (Döring et al., 2015).

Furthermore, while 2023 was marked by intense drought and heat stress, 2022 was equally affected (Fig. S1-S3), and 2021 on the contrary was struck by heavy rainfall throughout the spring/summer and hail events during grain formation and ripening. This highlights the fact that every year or environment brings its own kind of stressors, which can also cumulate – increasingly so with global warming and the expected rise in the frequency of intense weather events (Allan et al., 2021) – making it difficult to properly disentangle the effects of one stressor (Herrera et al., 2020). For instance, in our study it is impossible to properly assess the effects of intense heat, since our “cooler” year of comparison was affected by extreme rainfall. To properly disentangle the effects of weather, we would either need more experiments across more environmental conditions to cover a larger gradient of stressors, or it would require to experimentally manipulate the stressors, through rainout shelters or irrigation for instance (Jiang et al., 2020; Wright et al., 2021; Yahdjian & Sala, 2002). In addition, measuring the microclimates in plant communities would also provide valuable further insights into mechanistic processes taking place in diverse systems, such as local increase in moisture or cooler temperatures below the canopy (Aguirre et al., 2021; Kemppinen et al., 2024). Direct measurements of plant stress levels (e.g. stomatal conductance) would additionally enable to precisely identify stress level and duration (Wang et al., 2015).

## Conclusions

This study showed that mixtures generally outperformed monocultures in terms of global performance and stability for 5 crop parameters reflecting grain yield and quality. Notably, for most parameters we observed an increased in stability in mixtures. We also found a positive effect of mixtures on Zeleny sedimentation value, showing the potential of variety mixtures to improve crop quality. Furthermore, our study allowed to emphasize the role of competition for light for mixture benefits, notably the role of height and leaf area index. A higher light interception indeed led to increased overyielding and overquality. Despite variability across environmental conditions, wheat varieties mixtures are thus generally worth it. However, the varying performances of the different mixtures underline the importance of a thorough assessment of the characteristics of the individual components that are used to design the mixtures. As a practical recommendation, we advise to combine varieties that have similar heights and phenologies but different tillering abilities and yield potentials, and highlight the importance to monitor light interception. We also provide a list of promising mixtures readily available for farmers. All in all, we show that variety mixtures represent a promising solution to maintain and increase wheat yield, quality, and stability; this study could be of interest to farmers, processors, and bakers and help promote the larger adoption of wheat variety mixtures.

## Acknowledgments

We thank Yann Imhoff, Noémie Schaad, Julie Roux, Reynold Bovy, Flavio Foiada and Malgorzata Watroba for their assistance with field experiments, and Patrick Krähenbühl for help with the sedimentation analyses. We also acknowledge the support from Michael Winzeler and Hans Winzeler regarding the choice of the accessions, and the interpretation of the results. This project was jointly funded by the Swiss Federal Office for Agriculture (BLW) and IP-Suisse.

## Author contributions

**Laura Stefan:** Investigation, Data Curation, Formal analysis, Project administration, Visualization, Writing – Original Draft, Writing – Review and Editing. **Silvan Strebel:** Investigation, Writing – Review and Editing. **Karl-Heinz Camp:** Conceptualization, Funding acquisition, Resources, Investigation, Writing – Review and Editing. **Sarah Christinat:** Conceptualization, Funding acquisition, Writing – Review and Editing. **Dario Fossati:** Conceptualization, Funding acquisition, Resources, Writing – Review and Editing. **Christian Städeli:** Conceptualization, Funding acquisition, Writing – Review and Editing. **Lilia Levy Häner:** Conceptualization, Funding acquisition, Resources, Investigation, Supervision, Writing – Review and Editing.

## Declaration of interests

The authors declare that they have no conflict of interest.

## Funding sources

This project was jointly funded by the Swiss Federal Office for Agriculture (BLW) and IP-Suisse.

## Data availability statement

The data will be available on Zenodo upon publication of the manuscript.

## Notes

### Competing Interest Statement

The authors have declared no competing interest.

