## Supplementary Material for "Multi trait assessment of wheat variety mixtures performance and stability: mixtures for the win!"

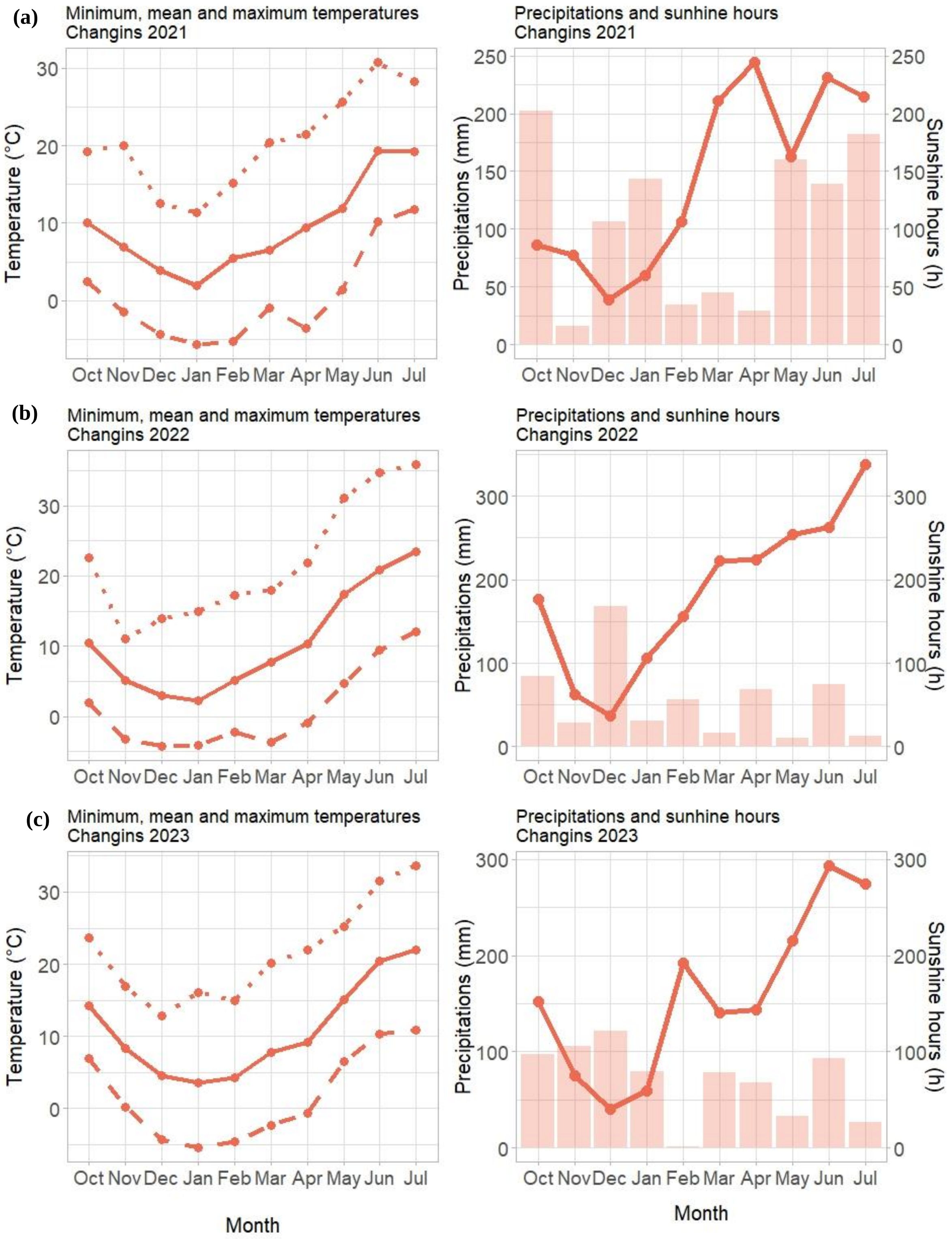


**Fig. S1: Left panel: Minimum, mean, and maximum temperatures in Changins 2021 (a), 2022 (b) and 2023 (c).** Dashed lines represent minimum temperatures, full lines represent the mean, and dotted lines represent maximum temperatures. **Right panel: Precipitations and sunshine hours.** Lines represent the sunshine hours, while bars show the precipitation.

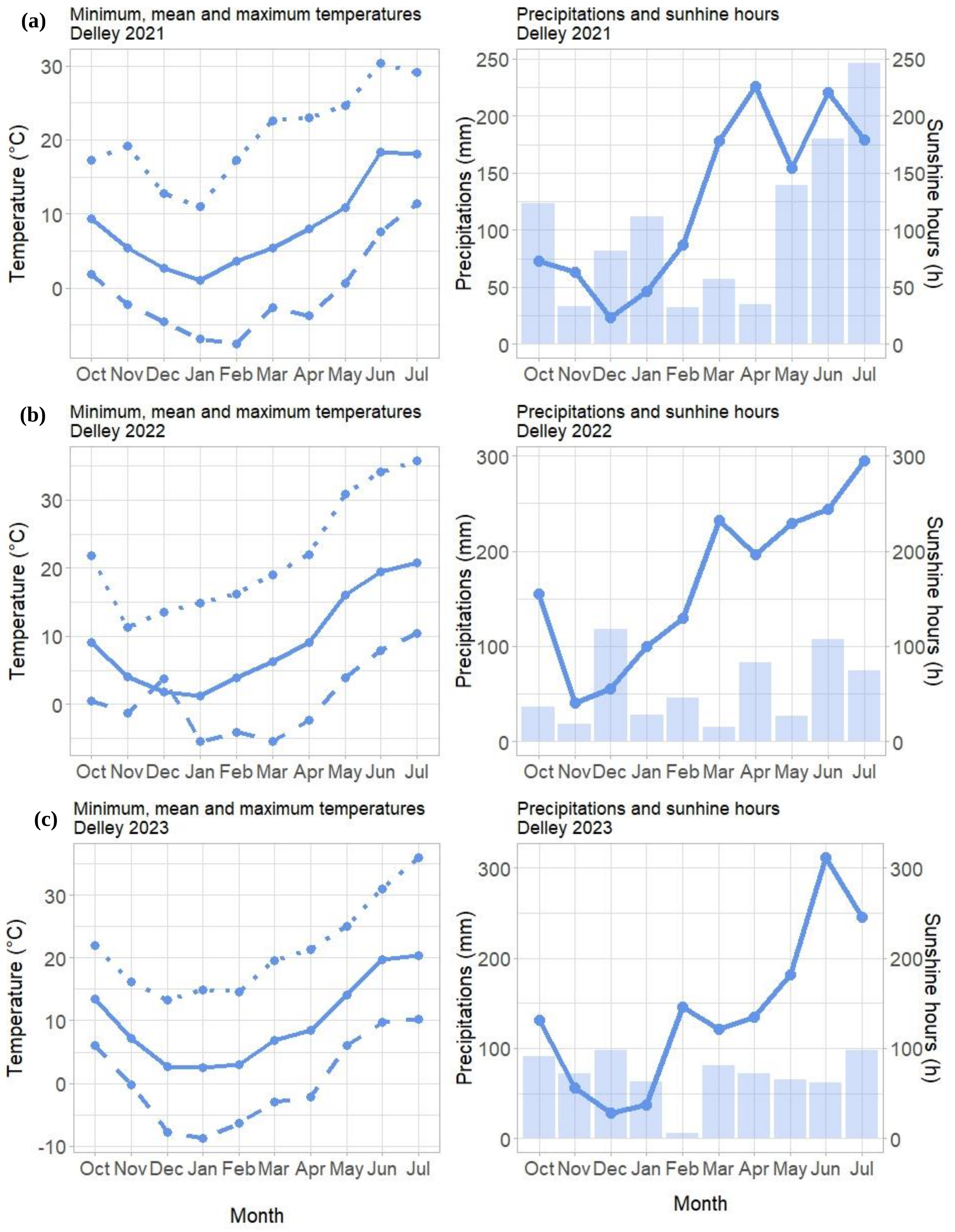


**Fig. S2: Left panel: Minimum, mean, and maximum temperatures in Delley 2021 (a), 2022 (b) and 2023 (c).** Dashed lines represent minimum temperatures, full lines represent the mean, and dotted lines represent maximum temperatures. **Right panel: Precipitations and sunshine hours.** Lines represent the sunshine hours, while bars show the precipitation.

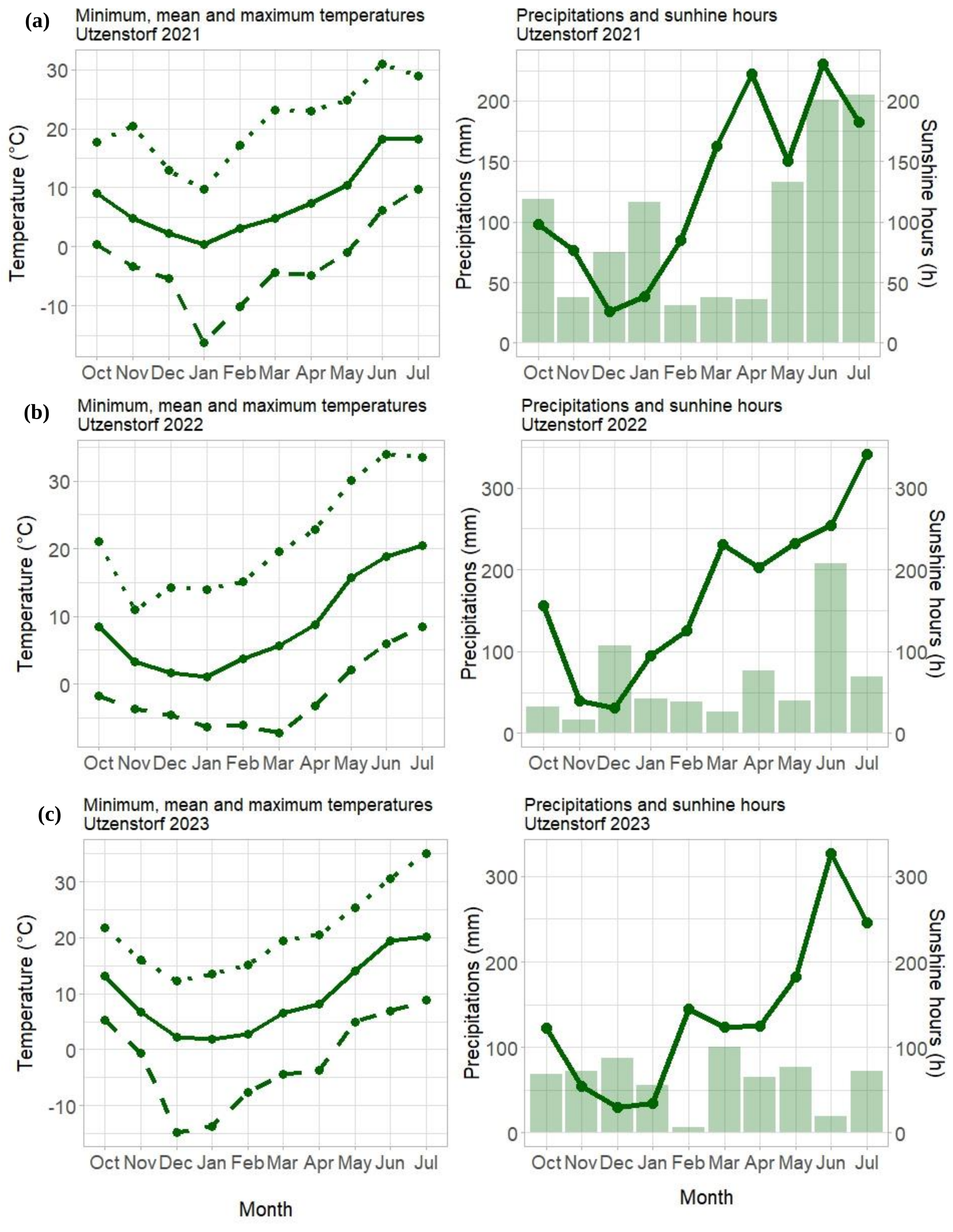


**Fig. S3: Left panel: Minimum, mean, and maximum temperatures in Utzenstorf 2021 (a), 2022 (b) and 2023 (c).** Dashed lines represent minimum temperatures, full lines represent the mean, and dotted lines represent maximum temperatures. **Right panel: Precipitations and sunshine hours.** Lines represent the sunshine hours, while bars show the precipitation.

**Table S1: Experimental details and soil status of the fields used for the experimental trials**

| | *Changins 2021* | *Changins 2022* | *Changins 2023* | *Delley 2021* | *Delley 2022* | *Delley 2023* | *Utzenstorf 2021* | *Utzenstorf 2022* | *Utzenstorf 2023* |
|---|---|---|---|---|---|---|---|---|---|
| *Sowing date* | 20.10.2020 | 13.10.2021 | 18.10.2022 | 16.10.2020 | 15.10.2021 | 18.10.2022 | 20.10.2020 | 28.10.2021 | 19.10.2022 |
| *Harvest date* | 23.07.2021 | 11.07.2022 | 11.07.2023 | 23.07.2021 | 13.07.2022 | 11.07.2023 | 23.07.2021 | 18.07.2022 | 17.07.2023 |
| *% Clay* | 26 | 22 | 21 | 14 | 20 | 15 | 15 – 20 | 15 – 20 | 15 – 20 |
| *% Silt* | 43 | 47 | 43 | 28 | 46 | NA | NA | NA | NA |
| *% Sand* | 31 | 31 | 36 | 58 | 34 | NA | NA | NA | NA |
| *% Organic Matter* | 2.9 | 2.5 | 2.2 | 1.5 | 2.2 | 1.5 | 2 – 4.9 | 2 – 4.9 | 2 – 4.9 |
| *pH* | 7.7 | 7.6 | 7.1 | 6.9 | 8 | 6.3 | 6.4 | 5.7 | 6.9 |

**Table S2: Description of the accessions/varieties used for the experimental trials.**
Data presented here originate from the national variety testing trials and were averaged over years and sites.

| Experimental number | Name | Quality Class | Presence of awns | Standardized Yield (dt/ha) | Zeleny sedimentation rate (ml) | Protein content (%) | Relative heading date | Height (cm) |
|---|---|---|---|---|---|---|---|---|
| 111.15885 | FALOTTA | 1 | Awns | 104.6 | 56.5 | 14.0 | 1.8 | 88.1 |
| 111.16373 | | NA | No awns | 93.2 | 54.3 | 14.4 | 0.4 | 88.2 |
| 111.14470 | COLMETTA | 2 | Awns | 113.0 | 46.5 | 11.9 | -1.4 | 87.5 |
| 111.15797 | CAMPANILE | 1 | No awns | 112.0 | 52.8 | 12.9 | 1.2 | 94.4 |
| 111.15874 | SCHILTHORN | TOP | No awns | 107.0 | 60.9 | 13.5 | -0.1 | 91.8 |
| 211.14074 | | NA | Awns | 107.0 | 68.5 | 12.7 | -2.4 | **100.7** |
| 111.15974 | BODELI | TOP | Awns | 100.8 | 62.5 | 14.1 | -0.6 | 96.6 |
| 111.13431 | MOLINERA | TOP | Awns | 89.3 | 62.5 | 15.0 | 0.2 | 87.8 |

**Table S3**. **Type-I Analysis of Variance Table of Yield (dt/ha), Protein content (%), Thousand kernel weight (TKW, g), Hectoliter weight (kg/hl), and Zeleny sedimentation value (ml) in response to experimental factors (year, site) and diversity treatment (mono vs. mix, variety number).**

*DenDF*, degrees of freedom of error term; *NumDF*, degrees of freedom of term; *F-value*, variance ratio; *Pr(>F)*, error probability. P-values in bold are significant at α = 0.05; * (P < 0.05), ** (P < 0.01), *** (P < 0.001). n = 996

| | | *Yield* | *Yield* | *Protein* | *Protein* | *TKW* | *TKW* | *HLW* | *HLW* | *Zeleny* | *Zeleny* |
|---|---|---|---|---|---|---|---|---|---|---|---|
| | *NumDF* | *F value* | *Pr(>F)* | *F value* | *Pr(>F)* | *F value* | *Pr(>F)* | *F value* | *Pr(>F)* | *F value* | *Pr(>F)* |
| *Year* | 2 | 39.93 | **<0.001*** | 9.71 | **0.0013** | 5.77 | **0.015*** | 15.36 | **<0.001*** | 0.94 | 0.41 |
| *Site* | 2 | 80.29 | **<0.001*** | 9.30 | **0.0016** | 31.84 | **<0.001*** | 1.16 | 0.33 | 14.65 | **<0.001*** |
| *Mono vs. mix* | 1 | 0.0016 | 0.96 | 0.12 | 0.72 | 0.001 | 0.97 | 0.30 | 0.58 | 5.18 | **0.022*** |
| *Variety number* | 1 | 0.0005 | 0.98 | 0.072 | 0.78 | 0.02 | 0.86 | 0.71 | 0.39 | 0.074 | 0.78 |
| *Year x Site* | 4 | 44.77 | **<0.001*** | 0.48 | 0.74 | 40.21 | **<0.001*** | 6.19 | **0.0026** | 2.58 | 0.072 |
| *Year x Mono vs. mix* | 2 | 0.30 | 0.74 | 0.19 | 0.82 | 1.65 | 0.19 | 0.44 | 0.63 | 1.66 | 0.18 |
| *Year x Var. number* | 2 | 1.09 | 0.33 | 0.55 | 0.57 | 1.15 | 0.31 | 1.08 | 0.33 | 0.35 | 0.70 |
| *Site x Mono vs. mix* | 2 | 2.73 | 0.065 | 1.93 | 0.14 | 0.10 | 0.90 | 0.77 | 0.45 | 1.65 | 0.19 |
| *Site x Var. number* | 2 | 0.55 | 0.57 | 2.14 | 0.11 | 0.65 | 0.52 | 2.97 | 0.051 | 2.28 | 0.10 |
| *Year x Site x Mono vs. mix* | 4 | 0.98 | 0.41 | 1.40 | 0.23 | 0.33 | 0.85 | 0.18 | 0.94 | 0.25 | 0.90 |
| *Year x Site x Var. number* | 4 | 2.61 | **0.034*** | 2.21 | 0.065 | 0.63 | 0.63 | 1.91 | 0.11 | 0.99 | 0.41 |

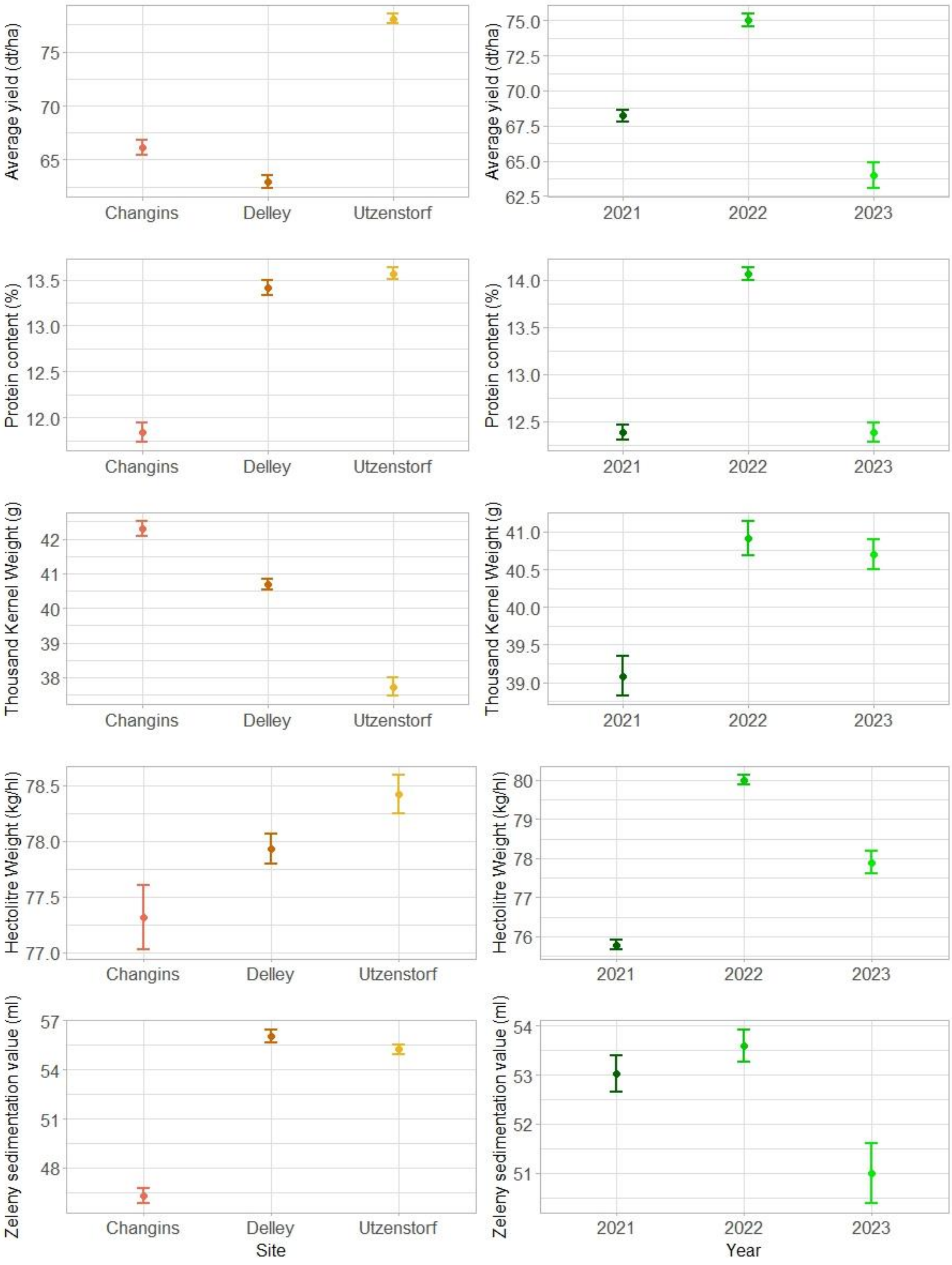


**Fig. S4: Performance of wheat communities averaged across sites (left panel) and years (right panel).**

Dots represent the mean values across plots; lines represent the standard error. n=996

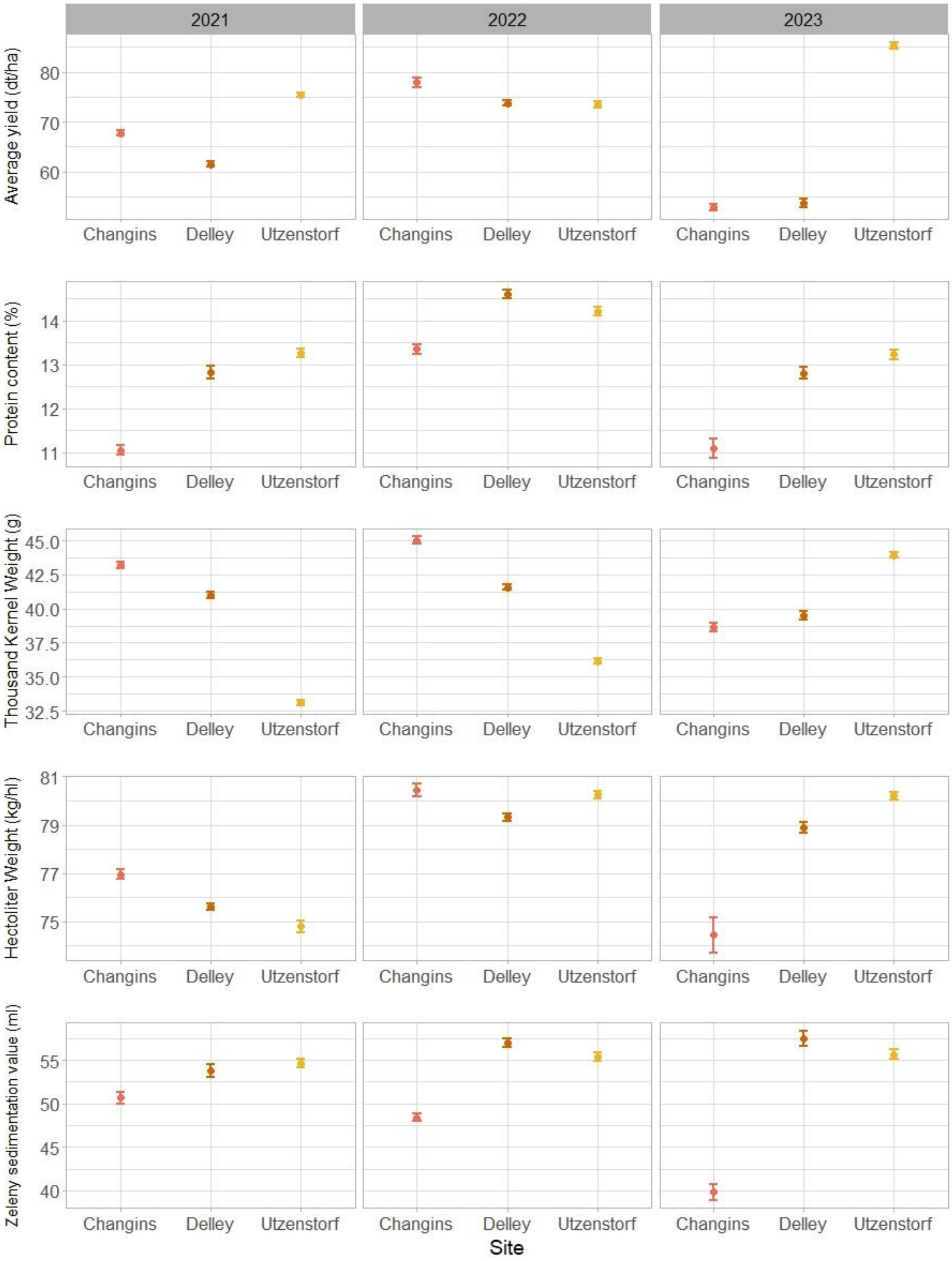


**Fig. S5: Performance of wheat communities across environments.**
Dots represent the mean values across plots; lines represent the standard error. n=996

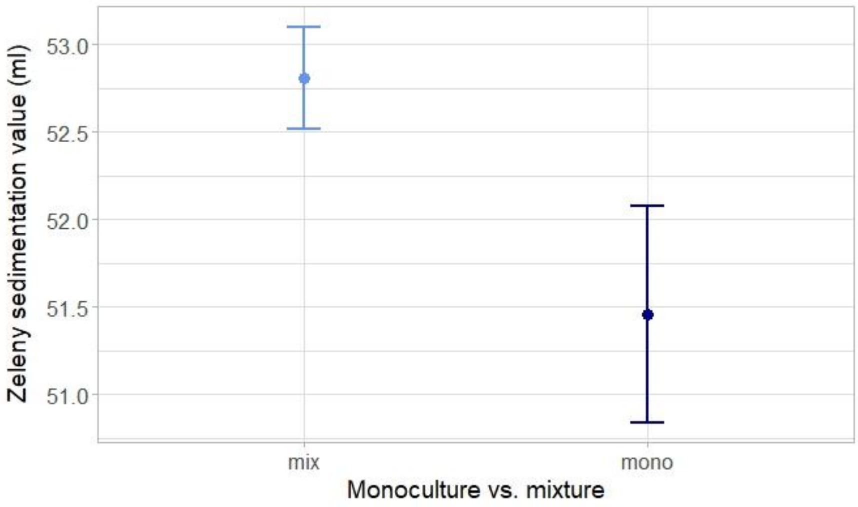


**Fig. S6: Zeleny sedimentation value (ml) in response to diversity treatment (monoculture vs. mixture) across environments.**

Dots represent the mean values across plots; lines represent the standard error. n=996

**Table S4**. **Type-I Analysis of Variance Table of Overperformance (overyield (dt/ha), overprotein content (%), overTKW (g), overHLW (kg/hl), and overZeleny (ml)) in response to experimental factors (year, site) and diversity treatment (variety number).**

*DenDF*, degrees of freedom of error term; *NumDF*, degrees of freedom of term; *F-value*, variance ratio; *Pr(>F)*, error probability. P-values in bold are significant at α = 0.05; * (P < 0.05), ** (P < 0.01), *** (P < 0.001). n = 780

| | *NumDF* | *Overyield F value* | *Overyield Pr(>F)* | *Overprotein F value* | *Overprotein Pr(>F)* | *OverTKW F value* | *OverTKW Pr(>F)* | *OverHLW F value* | *OverHLW Pr(>F)* | *OverZeleny F value* | *OverZeleny Pr(>F)* |
|---|---|---|---|---|---|---|---|---|---|---|---|
| *Year* | 2 | 1.03 | 0.35 | 0.67 | 0.51 | 8.08 | **<0.001***** | 1.96 | 0.14 | 11.79 | **<0.001***** |
| *Site* | 2 | 9.67 | **<0.001***** | 7.05 | **<0.001***** | 0.58 | 0.55 | 3.36 | **0.035*** | 11.93 | **<0.001***** |
| *Variety number* | 1 | 0.0033 | 0.95 | 1.2 | 0.27 | 0.54 | 0.46 | 0.51 | 0.48 | 0.10 | 0.74 |
| *Year x Site* | 4 | 3.44 | **0.0083**** | 5.12 | **<0.001***** | 1.55 | 0.18 | 0.81 | 0.51 | 1.77 | 0.13 |
| *Year x Var. number* | 2 | 0.81 | 0.44 | 0.44 | 0.64 | 1.25 | 0.28 | 1.06 | 0.34 | 0.52 | 0.58 |
| *Site x Var. number* | 2 | 0.42 | 0.65 | 1.71 | 0.18 | 0.74 | 0.47 | 2.94 | 0.053 | 3.45 | **0.032*** |
| *Year x Site x Var. number* | 4 | 1.94 | 0.10 | 1.78 | 0.13 | 0.72 | 0.57 | 1.87 | 0.11 | 1.49 | 0.20 |

**Table S5.** **Results of the t-test to evaluate whether overperformance is significantly different from 0.** Numbers indicate the average values per environment. Bold numbers indicate environments where the t-test was significant.

| *t.test* | *Changins.2021* | *Changins.2022* | *Changins.2023* | *Delley.2021* | *Delley.2022* | *Delley.2023* | *Utzenstorf.2021* | *Utzenstorf.2022* | *Utzenstorf.2023* |
|---|---|---|---|---|---|---|---|---|---|
| *Overyield* | **1.72** | **2.8** | 0.65 | 0.49 | **-1.7** | -1.6 | -0.43 | **-2.14** | 0.93 |
| *Overprotein* | **-0.25** | **0.35** | -0.033 | -0.13 | **-0.41** | **-0.36** | -0.14 | -0.14 | 0.084 |
| *OverTKW* | **-0.49** | **0.42** | 0.29 | -0.19 | **0.29** | 0.10 | -0.19 | -0.05 | 0.07 |
| *OverHLW* | 0.004 | **-0.79** | -0.88 | -0.05 | **0.31** | -0.21 | 0.28 | **0.28** | **-0.27** |
| *OverZeleny* | **-0.74** | 0.49 | -0.21 | 0.84 | **2.23** | 1.5 | -0.006 | **3.8** | **1.8** |
| *Overdensity* | **24.5** | **37.3** | 3.7 | **-28.1** | -6.8 | -4 | **-30.5** | -6.3 | 2 |

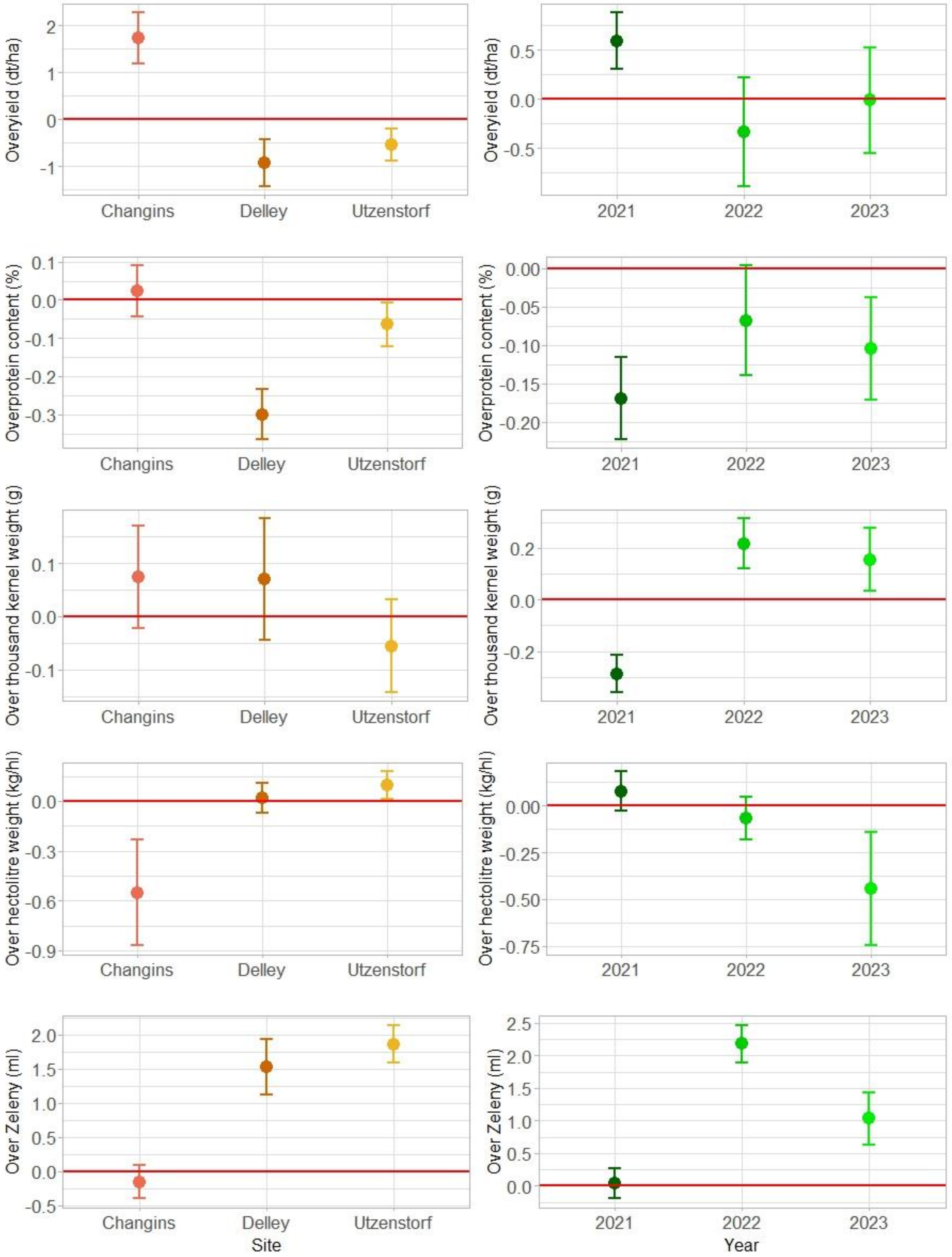


**Fig. S7: Overperformance averaged across sites (left panel) and years (right panel).** Dots represent the mean values across plots; lines represent the standard error. n=780

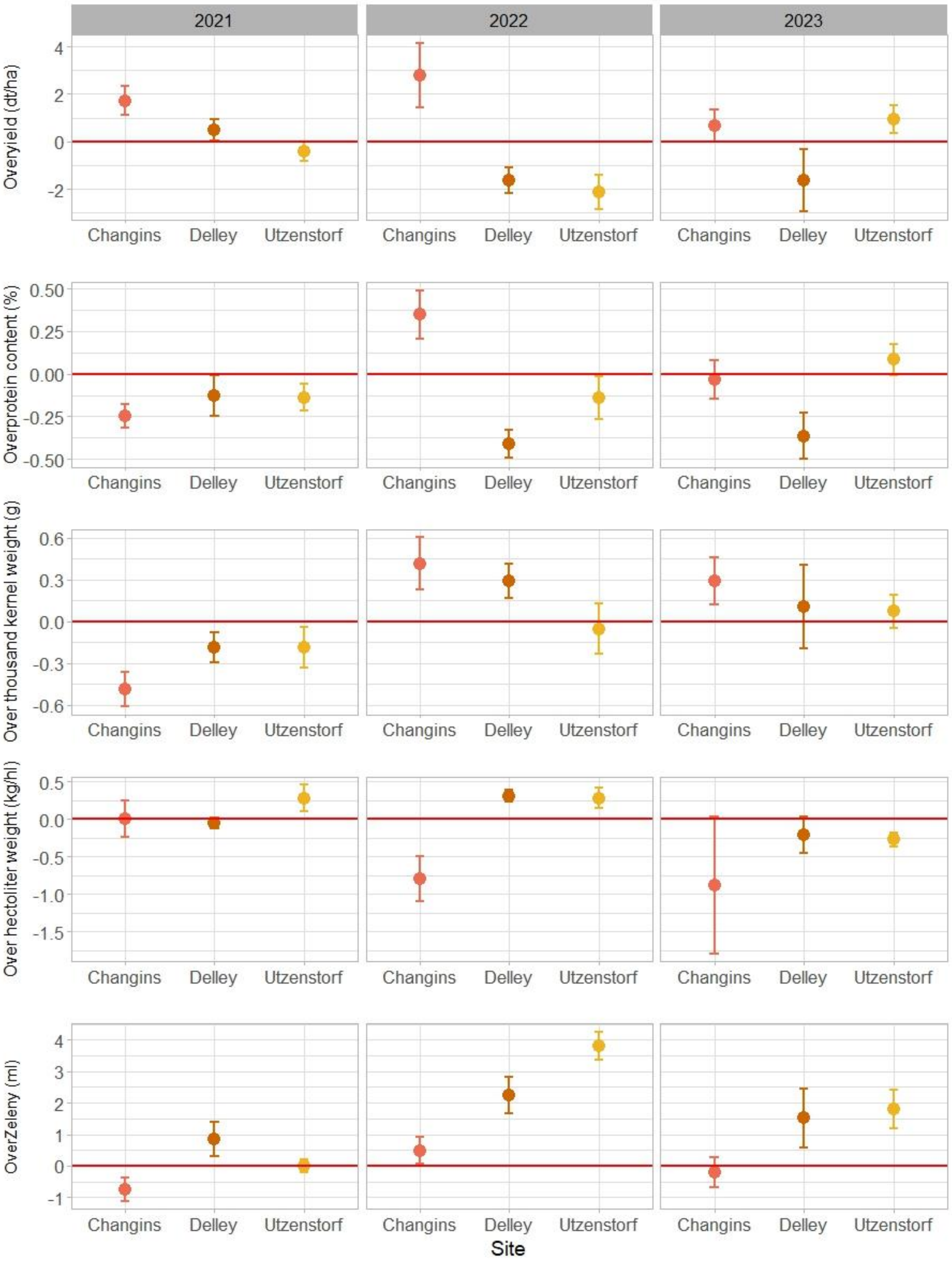


**Fig. S8: Overperformance in response to environment.**
Dots represent the mean values across plots; lines represent the standard error. n=780

**Table S6**. **Type-III Analysis of Variance Table of Overperformance (overyield (dt/ha), overprotein content (%), overTKW (g), overHLW (kg/hl), and overZeleny (mL)) in response to explanatory variables from varieties in monocultures.**

*DenDF*, degrees of freedom of error term; *NumDF*, degrees of freedom of term; *F-value*, variance ratio; *Pr(>F)*, error probability. P-values in bold are significant at α = 0.05; * (P < 0.05), ** (P < 0.01), *** (P < 0.001). n = 752

| | | *Overyield* | *Overyield* | *Overprotein* | *Overprotein* | *OverTKW* | *OverTKW* | *OverHLW* | *OverHLW* | *OverZeleny* | *OverZeleny* |
|---|---|---|---|---|---|---|---|---|---|---|---|
| | *NumDF* | *F value* | *Pr(>F)* | *F value* | *Pr(>F)* | *F value* | *Pr(>F)* | *F value* | *Pr(>F)* | *F value* | *Pr(>F)* |
| *Awns difference* | 1 | 0.1383 | 0.712 | 0.04 | 0.84179 | 0.2172 | 0.644 | 0.2072 | 0.65199 | 0.5878 | 0.44351 |
| *Diff in mono yield* | 1 | 0.0522 | 0.8194 | 3.728 | **0.05397** | 0.304 | 0.58166 | 0.174 | 0.67678 | 6.62 | **0.01028*** |
| *Diff in mono protein* | 1 | 0.6796 | 0.4105 | 7.1143 | **0.00794**** | 0.3103 | 0.57799 | 0.0008 | 0.97764 | 2.3444 | 0.12617 |
| *Diff in mono height* | 1 | 16.7542 | **<0.001***** | 0.072 | 0.78866 | 0.2068 | 0.64976 | 6.6292 | **0.01082*** | 1.5666 | 0.2111 |
| *Diff in mono heading day* | 1 | 1.5029 | 0.2217 | 0.0295 | 0.86383 | 4.3291 | **0.03889*** | 0.0158 | 0.90002 | 3.4775 | **0.06262** |
| *Diff in mono density* | 1 | 0.0005 | 0.982 | 5.7392 | **0.01684*** | 0.8283 | 0.36309 | 1.8331 | 0.17625 | 1.7381 | 0.18781 |

**Table S7**. **Type-III Analysis of Variance Table of Overperformance in Changins for the three years, in response to explanatory variables from varieties in monocultures.**

*DenDF*, degrees of freedom of error term; *NumDF*, degrees of freedom of term; *F-value*, variance ratio; *Pr(>F)*, error probability. P-values in bold are significant at α = 0.05; * (P < 0.05), ** (P < 0.01), *** (P < 0.001). n = 248

| | | *Overyield* | *Overyield* | *Overprotein* | *Overprotein* | *OverTKW* | *OverTKW* | *OverHLW* | *OverHLW* | *OverZeleny* | *OverZeleny* |
|---|---|---|---|---|---|---|---|---|---|---|---|
| | *NumDF* | *F value* | *Pr(>F)* | *F value* | *Pr(>F)* | *F value* | *Pr(>F)* | *F value* | *Pr(>F)* | *F value* | *Pr(>F)* |
| *Diff in mono LAI early* | 1 | 2.2876 | 0.1321 | 0.2799 | 0.5975 | 2.6199 | 0.107587 | 0.1637 | 0.6862 | 1.9294 | 0.167183 |
| *Diff in mono LAI late* | 1 | 0.397 | 0.5295 | 0.7379 | 0.3917 | 0.3202 | 0.572644 | 0.4114 | 0.5222 | 0.0754 | 0.784039 |
| *OverLAI early* | 1 | 0.5013 | 0.4798 | 23.6146 | **<0.001***** | 0.2424 | 0.623051 | 0.489 | 0.4853 | 7.9098 | **0.0054**** |
| *OverLAI late* | 1 | 25.9801 | **<0.001***** | 0.0384 | 0.8448 | 10.4175 | **0.001475**** | 0.0241 | 0.8767 | 2.6434 | 0.105658 |

**Table S8**. **Type-III Analysis of Variance Table of Overperformance in Changins in 2022 and 2023, in response to explanatory variables from varieties in monocultures.**
*DenDF*, degrees of freedom of error term; *NumDF*, degrees of freedom of term; *F-value*, variance ratio; *Pr(>F)*, error probability. P-values in bold are significant at α = 0.05; * (P < 0.05), ** (P < 0.01), *** (P < 0.001). n = 165

| | | *Overyield* | *Overyield* | *Overprotein* | *Overprotein* | *OverTKW* | *OverTKW* | *OverHLW* | *OverHLW* | *OverZeleny* | *OverZeleny* |
|---|---|---|---|---|---|---|---|---|---|---|---|
| | *NumDF* | *F value* | *Pr(>F)* | *F value* | *Pr(>F)* | *F value* | *Pr(>F)* | *F value* | *Pr(>F)* | *F value* | *Pr(>F)* |
| *Diff.SLA* | 1 | 0.59 | 0.44 | 1.48 | 0.22 | 0.64 | 0.43 | 0.0003 | 0.99 | 0.55 | 0.46 |
| *Diff.LDMC* | 1 | 1.4 | 0.24 | 4.34 | **0.039*** | 0.0037 | 0.95 | 1.95 | 0.17 | 1.65 | 0.20 |

**Table S9**. **Type-I Analysis of Variance Table of Stability per site (temporal stability) for yield, protein content, TKW, HLW, and Zeleny, as well as MTSI, in response to experimental factors (site) and diversity treatment (mono vs. mix and variety number).**
*DenDF*, degrees of freedom of error term; *NumDF*, degrees of freedom of term; *F-value*, variance ratio; *Pr(>F)*, error probability. P-values in bold are significant at α = 0.05; * (P < 0.05), ** (P < 0.01), *** (P < 0.001). n = 111

| | | *WAASB Yield* | *WAASB Yield* | *WAASB Protein* | *WAASB Protein* | *WAASB TKW* | *WAASB TKW* | *WAASB HLW* | *WAASB HLW* | *WAASB Zeleny* | *WAASB Zeleny* | *MTSI* | *MTSI* |
|---|---|---|---|---|---|---|---|---|---|---|---|---|---|
| | *NumDF* | *F value* | *Pr(>F)* | *F value* | *Pr(>F)* | *F value* | *Pr(>F)* | *F value* | *Pr(>F)* | *F value* | *Pr(>F)* | *F value* | *Pr(>F)* |
| *Site* | 2 | 0.9553 | 0.38794 | 9.2217 | **0.000201***** | 4.7757 | **0.01128*** | 5.1916 | **0.00783**** | 3.5875 | **0.032811*** | 71.5718 | **<0.001***** |
| *Mono vs. mix* | 1 | 0.3369 | 0.56286 | 4.3292 | **0.039829*** | 7.3672 | **0.01013*** | 2.1379 | 0.15237 | 9.4344 | **0.004108**** | 4.6628 | **0.03756*** |
| *Variety number* | 2 | 1.7213 | 0.18375 | 0.4368 | 0.64726 | 1.3471 | 0.27279 | 0.017 | 0.98315 | 0.2464 | 0.782923 | 1.2952 | 0.2863 |
| *Site x Mono vs. mix* | 2 | 2.551 | 0.082 | 1.9036 | 0.153993 | 0.1183 | 0.88857 | 3.3269 | **0.0415*** | 1.9192 | 0.154312 | 0.6113 | 0.54546 |
| *Site x Var. number* | 4 | 0.7155 | 0.58315 | 1.9159 | 0.112958 | 0.1813 | 0.94738 | 1.052 | 0.38665 | 0.412 | 0.799436 | 0.9527 | 0.43882 |

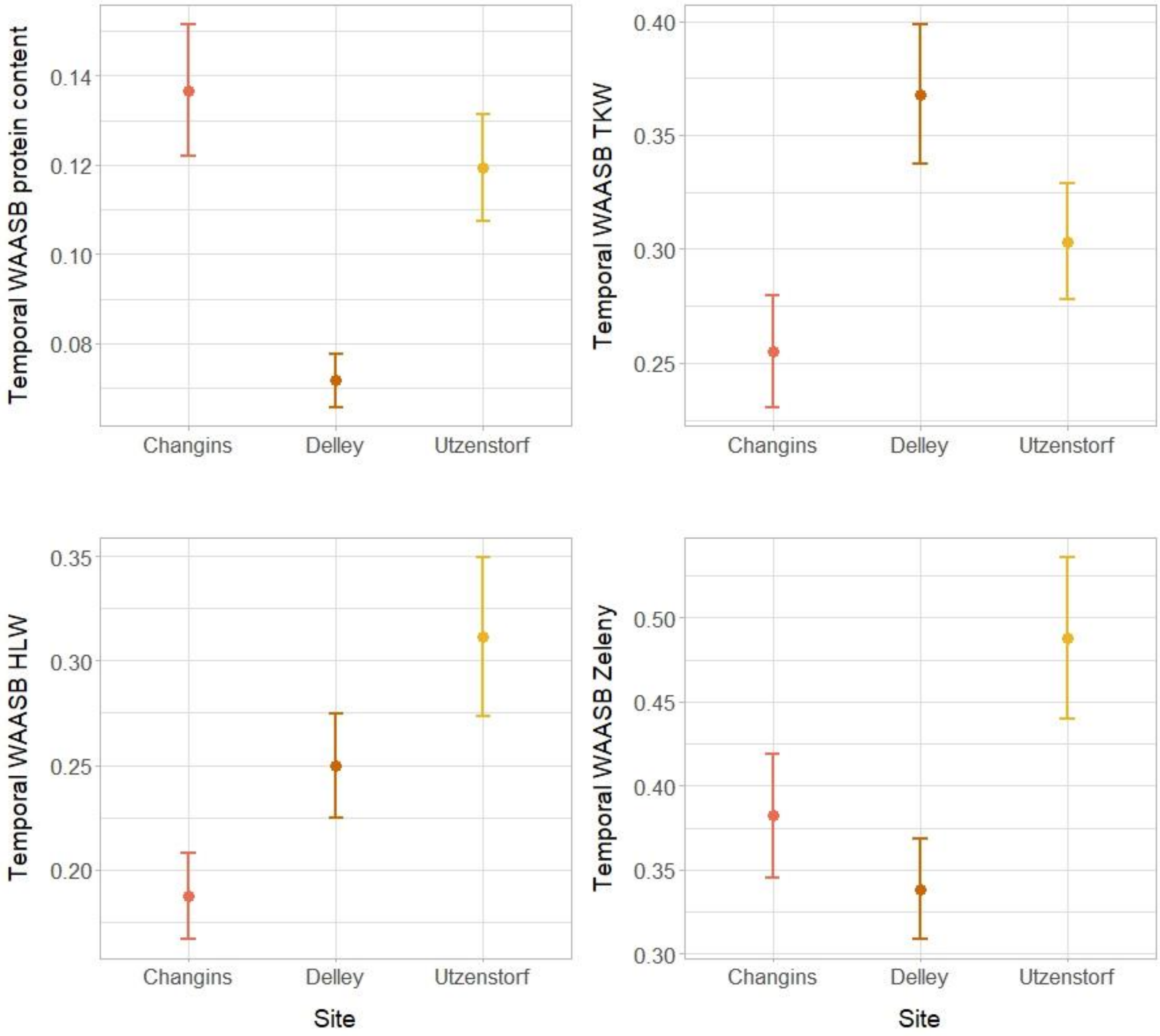


**Fig. S9: Temporal Stability of protein content, TKW, HLW and Zeleny in response to site.**

Dots represent the mean values across plots; lines represent the standard error. n=111

**Table S10**. **Type-III Analysis of Variance Table of Stability per site (temporal stability) for yield, protein content, TKW, HLW, and Zeleny, as well as MTSI, in response to explanatory variables from varieties in monocultures.**

*DenDF*, degrees of freedom of error term; *NumDF*, degrees of freedom of term; *F-value*, variance ratio; *Pr(>F)*, error probability. P-values in bold are significant at α = 0.05; * (P < 0.05), ** (P < 0.01), *** (P < 0.001). n = 84

| | | *WAASB Yield* | *WAASB Yield* | *WAASB Protein* | *WAASB Protein* | *WAASB TKW* | *WAASB TKW* | *WAASB HLW* | *WAASB HLW* | *WAASB Zeleny* | *WAASB Zeleny* | *MTSI* | *MTSI* |
|---|---|---|---|---|---|---|---|---|---|---|---|---|---|
| | *NumDF* | *F value* | *Pr(>F)* | *F value* | *Pr(>F)* | *F value* | *Pr(>F)* | *F value* | *Pr(>F)* | *F value* | *Pr(>F)* | *F value* | *Pr(>F)* |
| *Awns difference* | 1 | 0.0035 | 0.95287 | 4.8492 | **0.03073*** | 2.4742 | 0.1199 | 0.2897 | 0.5959 | 0.0112 | 0.9166 | 1.1322 | 0.2979 |
| *Diff in mono yield* | 1 | 0.1974 | 0.65809 | 0.5178 | 0.47399 | 0.2488 | 0.6193 | 0.0153 | 0.902 | 0.1685 | 0.6827 | 0.1409 | 0.7085 |
| *Diff in mono protein* | 1 | 1.7583 | 0.18876 | 0.1489 | 0.70072 | 0.4329 | 0.5126 | 0.3017 | 0.5855 | 0.6498 | 0.4239 | 1.619 | 0.2089 |
| *Diff in mono height* | 1 | 0.412 | 0.52288 | 0.0708 | 0.79091 | 1.4701 | 0.2291 | 0.23 | 0.6341 | 0.0007 | 0.9793 | 2.031 | 0.1607 |
| *Diff in mono heading day* | 1 | 2.8707 | 0.09425 | 0.0108 | 0.91746 | 0.3069 | 0.5812 | 0.1551 | 0.6961 | 1.5237 | 0.2234 | 0.0231 | 0.8799 |
| *Diff in mono density* | 1 | 2.288 | 0.13447 | 0.0818 | 0.77567 | 0.1022 | 0.75 | 1.9288 | 0.17 | 0.0825 | 0.7748 | 1.6398 | 0.2044 |

**Table S11**. **Type-III Analysis of Variance Table of temporal stability in Changins for yield, protein content, TKW, HLW, and Zeleny, as well as MTSI, in response to explanatory variables from varieties in monocultures.**

*DenDF*, degrees of freedom of error term; *NumDF*, degrees of freedom of term; *F-value*, variance ratio; *Pr(>F)*, error probability. P-values in bold are significant at α = 0.05; * (P < 0.05), ** (P < 0.01), *** (P < 0.001). n = 28

| | | *WAASB Yield* | *WAASB Yield* | *WAASB Protein* | *WAASB Protein* | *WAASB TKW* | *WAASB TKW* | *WAASB HLW* | *WAASB HLW* | *WAASB Zeleny* | *WAASB Zeleny* | *MTSI* | *MTSI* |
|---|---|---|---|---|---|---|---|---|---|---|---|---|---|
| | *NumDF* | *F value* | *Pr(>F)* | *F value* | *Pr(>F)* | *F value* | *Pr(>F)* | *F value* | *Pr(>F)* | *F value* | *Pr(>F)* | *F value* | *Pr(>F)* |
| *Diff in mono LAI early* | 1 | 0.8779 | 0.3585 | 0.5913 | 0.4497 | 1.3109 | 0.264 | 0.0023 | 0.96243 | 0.0315 | 0.8607 | 1.5056 | 0.2322 |
| *Diff in mono LAI late* | 1 | 1.4981 | 0.2334 | 1.0877 | 0.3078 | 0.0046 | 0.9467 | 3.4823 | 0.07483 | 0.0316 | 0.8606 | 0.0567 | 0.8139 |
| *Diff in SLA* | 1 | 0.5345 | 0.4721 | 0.0083 | 0.9282 | 0.1593 | 0.6935 | 0.1817 | 0.67391 | 0.2303 | 0.6359 | 1.3032 | 0.2654 |
| *Diff in LDMC* | 1 | 0.5435 | 0.4684 | 0.2395 | 0.6292 | 0.1636 | 0.6896 | 0.5235 | 0.47663 | 0.0636 | 0.8031 | 0.0485 | 0.8277 |

**Table S12**. **Type-I Analysis of Variance Table of Stability per year (spatial stability) for yield, protein content, TKW, HLW, and Zeleny, as well as MTSI, in response to experimental factors (year) and diversity treatment (mono vs. mix and variety number).**

*DenDF*, degrees of freedom of error term; *NumDF*, degrees of freedom of term; *F-value*, variance ratio; *Pr(>F)*, error probability. P-values in bold are significant at α = 0.05; * (P < 0.05), ** (P < 0.01), *** (P < 0.001). n = 111

| | | *WAASB Yield* | *WAASB Yield* | *WAASB Protein* | *WAASB Protein* | *WAASB TKW* | *WAASB TKW* | *WAASB HLW* | *WAASB HLW* | *WAASB Zeleny* | *WAASB Zeleny* | *MTSI* | *MTSI* |
|---|---|---|---|---|---|---|---|---|---|---|---|---|---|
| | *NumDF* | *F value* | *Pr(>F)* | *F value* | *Pr(>F)* | *F value* | *Pr(>F)* | *F value* | *Pr(>F)* | *F value* | *Pr(>F)* | *F value* | *Pr(>F)* |
| *Year* | 2 | 10.7907 | **<0.001*** ** | 26.9506 | **<0.001*** ** | 0.5694 | 0.56756 | 5.1913 | **0.007923** ** | 16.7616 | **<0.001*** ** | 100.6339 | **<0.001*** ** |
| *Mono vs. mix* | 1 | 0.0047 | 0.94567 | 0.5803 | 0.4512 | 4.6631 | **0.03303*** | 0.1825 | 0.671988 | 0.0001 | 0.9903 | 3.511 | 0.06909 |
| *Variety number* | 2 | 0.5259 | 0.59252 | 1.1509 | 0.3277 | 1.209 | 0.30251 | 0.1256 | 0.882348 | 0.4234 | 0.6581 | 0.5445 | 0.58486 |
| *Year x Mono vs. mix* | 2 | 3.5944 | **0.03084*** | 2.8997 | 0.0615 | 1.3666 | 0.25935 | 1.6712 | 0.195524 | 1.7012 | 0.1898 | 2.2948 | 0.10817 |
| *Year x Var. number* | 4 | 1.7614 | 0.14202 | 1.4523 | 0.2257 | 0.7135 | 0.58445 | 2.1609 | 0.082499 | 1.8072 | 0.137 | 3.056 | **0.02206*** |

**Table S13**. **Type-III Analysis of Variance Table of Stability per year (spatial stability) for yield, protein content, TKW, HLW, and Zeleny, as well as MTSI, in response to explanatory variables from varieties in monocultures.**

*DenDF*, degrees of freedom of error term; *NumDF*, degrees of freedom of term; *F-value*, variance ratio; *Pr(>F)*, error probability. P-values in bold are significant at α = 0.05; * (P < 0.05), ** (P < 0.01), *** (P < 0.001). n = 84

| | | *WAASB Yield* | *WAASB Yield* | *WAASB Protein* | *WAASB Protein* | *WAASB TKW* | *WAASB TKW* | *WAASB HLW* | *WAASB HLW* | *WAASB Zeleny* | *WAASB Zeleny* | *MTSI* | *MTSI* |
|---|---|---|---|---|---|---|---|---|---|---|---|---|---|
| | *NumDF* | *F value* | *Pr(>F)* | *F value* | *Pr(>F)* | *F value* | *Pr(>F)* | *F value* | *Pr(>F)* | *F value* | *Pr(>F)* | *F value* | *Pr(>F)* |
| *Awns difference* | 1 | 0.0052 | 0.9427 | 0.8744 | 0.3527 | 2.4812 | 0.11931 | 0.4847 | 0.4929 | 0.3509 | 0.5554 | 0.027 | 0.871 |
| *Diff in mono yield* | 1 | 2.4019 | 0.1254 | 0.0638 | 0.8013 | 1.0747 | 0.30314 | 1.164 | 0.2843 | 0.1701 | 0.6812 | 2.2146 | 0.1409 |
| *Diff in mono protein* | 1 | 0.1257 | 0.724 | 1.0616 | 0.3062 | 0.3403 | 0.56137 | 2.2534 | 0.138 | 0.0032 | 0.9552 | 0.4385 | 0.51 |
| *Diff in mono height* | 1 | 0.1361 | 0.7133 | 0.0151 | 0.9024 | 0.1001 | 0.75259 | 0.8091 | 0.3723 | 0.58 | 0.4487 | 0.2704 | 0.6048 |
| *Diff in mono heading day* | 1 | 2.1541 | 0.1463 | 0.1143 | 0.7362 | 0.4804 | 0.49032 | 0.948 | 0.3337 | 2.0064 | 0.1607 | 0.1033 | 0.7488 |
| *Diff in mono density* | 1 | 1.8468 | 0.1782 | 2.3994 | 0.1256 | 2.8677 | 0.09442 | 0.0311 | 0.8606 | 2.5656 | 0.1133 | 0.3657 | 0.5472 |

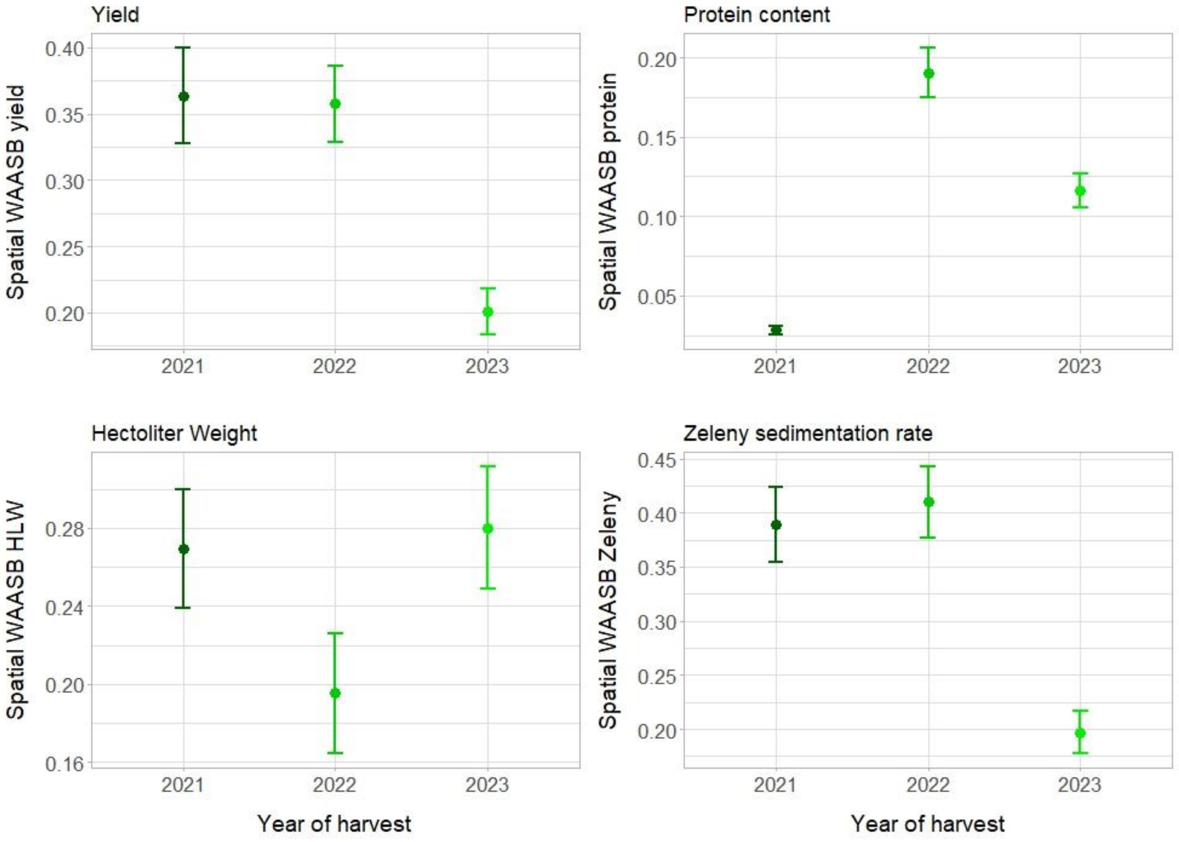


**Fig. S10: Spatial Stabiliy of yield, protein content, HLW and Zeleny, in response to year.** Dots represent the mean values across plots; lines represent the standard error. n=111

**Table S14**. **Type-I Analysis of Variance Table of overall Stability for yield, protein content, TKW, HLW, and Zeleny, as well as MTSI, in response to diversity treatment (mono vs. mix and variety number).**

*DenDF*, degrees of freedom of error term; *NumDF*, degrees of freedom of term; *F-value*, variance ratio; *Pr(>F)*, error probability. P-values in bold are significant at α = 0.05; * (P < 0.05), ** (P < 0.01), *** (P < 0.001). n = 37

| | | *WAASB Yield* | *WAASB Yield* | *WAASB Protein* | *WAASB Protein* | *WAASB TKW* | *WAASB TKW* | *WAASB HLW* | *WAASB HLW* | *WAASB Zeleny* | *WAASB Zeleny* | *MTSI* | *MTSI* |
|---|---|---|---|---|---|---|---|---|---|---|---|---|---|
| | *NumDF* | *F value* | *Pr(>F)* | *F value* | *Pr(>F)* | *F value* | *Pr(>F)* | *F value* | *Pr(>F)* | *F value* | *Pr(>F)* | *F value* | *Pr(>F)* |
| *Mono vs. mix* | 1 | 0.1125 | 0.7392 | 2.2209 | 0.1449 | 11.8389 | **0.001485**** | 0.0768 | 0.7833 | 4.6885 | **0.03707*** | 1.6198 | 0.2113 |
| *Variety number* | 2 | 0.4493 | 0.6416 | 0.1141 | 0.8925 | 1.6568 | 0.204962 | 0.7526 | 0.4784 | 0.777 | 0.46732 | 0.9946 | 0.3798 |

**Table S15**. **Type-III Analysis of Variance Table of overall Stability for yield, protein content, TKW, HLW, Zeleny, as well as MTSI, in response to explanatory variables from varieties in monocultures.**

*DenDF*, degrees of freedom of error term; *NumDF*, degrees of freedom of term; *F-value*, variance ratio; *Pr(>F)*, error probability. P-values in bold are significant at α = 0.05; * (P < 0.05), ** (P < 0.01), *** (P < 0.001). n = 28

| | | *WAASB Yield* | *WAASB Yield* | *WAASB Protein* | *WAASB Protein* | *WAASB TKW* | *WAASB TKW* | *WAASB HLW* | *WAASB HLW* | *WAASB Zeleny* | *WAASB Zeleny* | *MTSI* | *MTSI* |
|---|---|---|---|---|---|---|---|---|---|---|---|---|---|
| | *NumDF* | *F value* | *Pr(>F)* | *F value* | *Pr(>F)* | *F value* | *Pr(>F)* | *F value* | *Pr(>F)* | *F value* | *Pr(>F)* | *F value* | *Pr(>F)* |
| *Awns difference* | 1 | 0.1171 | 0.73558 | 1.7181 | 0.2041 | 0.025 | 0.87578 | 2.5738 | 0.12432 | 0.4758 | 0.49786 | 0.4216 | 0.523159 |
| *Diff in mono yield* | 1 | 3.3486 | 0.0815 | 0.1195 | 0.7331 | 2.8347 | 0.10706 | 2.0883 | 0.16392 | 0.2299 | 0.63658 | 10.81 | **0.003508**** |
| *Diff in mono protein* | 1 | 5.126 | **0.03428*** | 0.169 | 0.6852 | 3.154 | 0.09024 | 2.4751 | 0.13135 | 0.9984 | 0.32907 | 0.6505 | 0.428981 |
| *Diff in mono height* | 1 | 0.1055 | 0.74859 | 0.363 | 0.5533 | 11.0877 | **0.00318**** | 7.0549 | **0.01516*** | 0.4399 | 0.51438 | 7.5238 | **0.012189*** |
| *Diff in mono heading day* | 1 | 2.8751 | 0.10474 | 0.5247 | 0.4769 | 0.1402 | 0.71187 | 3.4444 | 0.07826 | 0.1234 | 0.72884 | 2.9244 | 0.101986 |
| *Diff in mono density* | 1 | 0.0243 | 0.87767 | 0.5167 | 0.4802 | 0.2053 | 0.65514 | 0.0209 | 0.88655 | 3.9176 | 0.06104 | 0.1251 | 0.727129 |

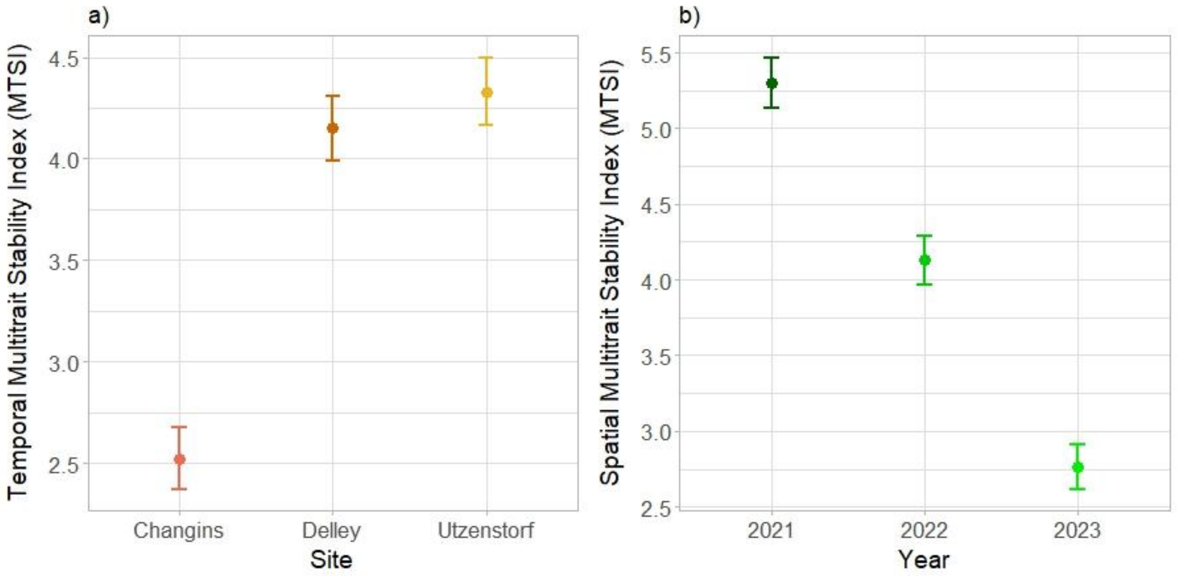


**Fig. S11: Temporal MTSI in response to site (a); Spatial MTSI in response to year (b)**
Dots represent the mean values across plots; lines represent the standard error. n=111

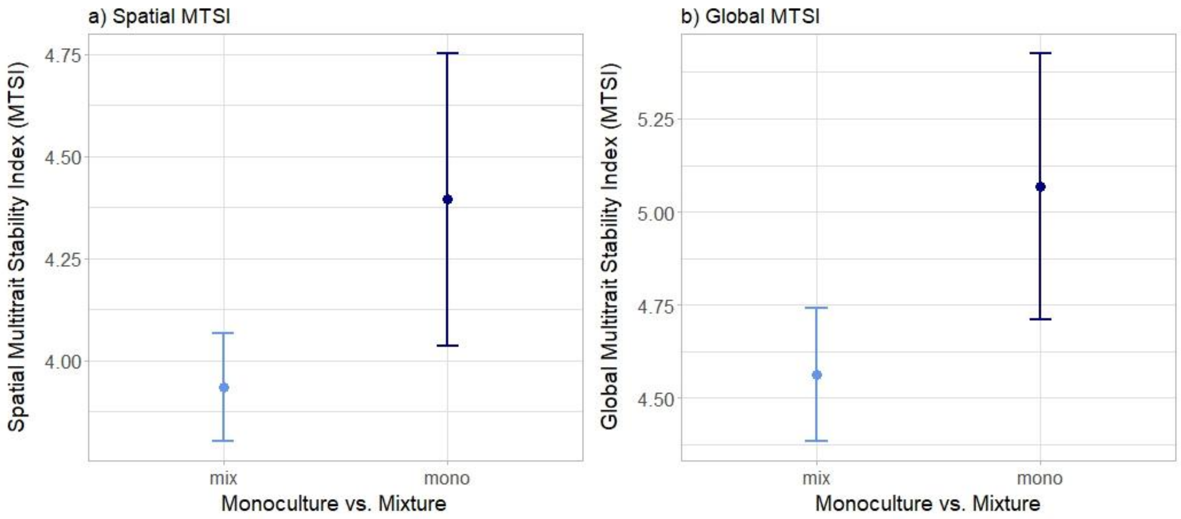


**Fig. S12: Spatial (a) and global (b) MTSI in response to diversity treatment (monoculture vs. mixture).**
Dots represent the mean values across plots; lines represent the standard error. n=111 (a); n=37 (b)
